# Genome-wide maps of nucleolus interactions reveal distinct layers of repressive chromatin domains

**DOI:** 10.1101/2020.11.17.386797

**Authors:** Cristiana Bersaglieri, Jelena Kresoja-Rakic, Shivani Gupta, Dominik Bär, Rostyslav Kuzyakiv, Raffaella Santoro

## Abstract

Eukaryotic chromosomes are folded into hierarchical domains, enabling the organization of the genome into functional compartments. Nuclear periphery and nucleolus are two nuclear landmarks thought to contribute to repressive chromosome architecture. However, while the role of nuclear lamina (NL) in genome organization has been well documented, the function of the nucleolus remains under-investigated due to the lack of methods for genome-wide maps of nucleolar associated domains (NADs). Here we established a method based on a Dam-fused engineered nucleolar histone H2B that marks DNA contacting the nucleolus. NAD-maps of ESCs and neural progenitors revealed layers of genome compartmentalization with distinct, repressive chromatin states based on the interaction with the nucleolus, NL, or both. NADs showed higher H3K9me2 and lower H3K27me3 content than regions exclusively interacting with NL. Upon ESC differentiation, chromosomes around the nucleolus acquire a more compact, rigid architecture whereas NADs specific for ESCs decrease their interaction strength within the repressive B-compartment strength, unlocking neural genes from repressive nuclear environment. The methodologies here developed will make possible to include the contribution of the nucleolus in future studies investigating the relationship between nuclear space and genome function.

## Introduction

In the nucleus of eukaryotic cells, chromosomes are arranged in a complex three-dimensional (3D) architecture that is thought to be important to ensure the correct execution of gene expression programs (Belmont, 2002; Dekker and Misteli, 2015; Kempfer and Pombo, 2019; Misteli, 2011; Nicodemi and Pombo, 2014). One important aspect of spatial genome organization is the local nuclear environment. Subnuclear compartments can serve as scaffold for chromatin tethering and allow the concentration of factors, thereby facilitating functions that rely on proteins found in limiting concentrations (Gonzalez-Sandoval and Gasser, 2016). Genomic interactions with the nuclear lamina (NL), which lies on the inner surface of the inner nuclear membrane, are mainly characterized by features typical of heterochromatin (van Steensel and Belmont, 2017). Lamina associated domains (LADs) contain genes in transcriptionally silent state or with low expression levels, have a low overall gene density, correspond to late-replicating DNA, and are typically enriched for repressive histone marks (Guelen et al., 2008; Peric-Hupkes et al., 2010; Pope et al., 2014). Another compartment suggested to serve as a scaffold for the location of repressive chromatin is the nucleolus, a membraneless sub-nuclear compartment that is the site of ribosome biogenesis (Bersaglieri and Santoro, 2019) and is made up of distinct, coexisting liquid phases (Feric et al., 2016). It has been suggested that the nucleolus and the NL might serve as interchangeable scaffolds for the localization of heterochromatic domains (Kind et al., 2013). However, the understanding of this genomic dynamics in the nuclear space remains still elusive. While the role of NL in genome organization has been well documented due to the identification and characterization of LADs in many cell types (van Steensel and Belmont, 2017), the function of the nucleolus remains under-investigated. One of the major reasons is that the identification of nucleolar associated domains (NADs) remains still a technical challenge. Previous attempts were based on the biochemical purification of nucleoli, a method that relies on sonication of nuclei, adjusting the power so that nucleoli remain intact while the rest of the nuclei are fragmented (Andersen et al., 2005; Desjardins et al., 1963; Sullivan et al., 2001). Using this methodology, first insights were provided into the composition of NADs, which appear to mainly consist of inactive regions (Dillinger et al., 2017; Nemeth et al., 2010; van Koningsbruggen et al., 2010). However, this method presents several technical limitations. First, since the heterochromatin is generally resistant to sonication (Becker et al., 2017), the identification of NADs upon sequencing of nucleoli purified through sonication can be biased toward repressive chromatin domains. Secondly, the experimental procedures to isolate nucleoli can be subjected to a certain variation, generating highly divergent NAD maps, even from the same cell types (Bizhanova et al., 2020; Lu et al., 2020). Third, it is difficult to achieve the purification of nucleoli in cells with open genome, such as embryonic stem cells (ESCs), unless protein-DNA crosslinking reagents are used, with a consequent extension of sonication time. Consequently, while data of LADs are frequently used in studies aimed to analyze genome organization in the cell’s nucleus, genomic contacts with the nucleolus have so far been excluded from these analyses. However, to fully understand the relationship between 3D genome organization and function, it is necessary to integrate also information on the organization of chromosomes around the nucleolus. This need calls for the establishment of novel methods for accurate genome-wide mapping of NADs that should be based on experimental procedures alternative to the highly variable biochemical isolation of nucleoli.

In this study, we established Nucleolar-DamID and HiC-rDNA methods that provided accurate genome-wide maps of NADs. The data revealed unprecedented layers of genome compartmentalization by showing distinct, repressive transcriptional and chromatin states based on the interaction with the nucleolus, NL, or both. NADs demarcate regions of the genome with a repressive state and are enriched in H3K9me2 and depleted in H3K27me3 relative to sequence only contacting NL. In ESCs, the chromatin state of ribosomal (r)RNA genes, which are located within the nucleolus, regulates the deposition of H3K9me2 at sequences adjacent to the nucleolus, indicating that nucleolus is not only a scaffold where repressive chromatin can be positioned but also it is part of the regulatory process for the establishment or maintenance of repressive chromatin states. Further, we showed that chromosome organization around the nucleolus changes according to developmental stage. Upon differentiation of ESCs into neural progenitors (NPCs), the architecture of all chromosomes surrounding the nucleolus becomes more rigid and compact, increasing the interaction frequency with centromere-proximal regions. Loss of nucleolar contacts during ESC differentiation coincides with an increase in the interaction strength within the active A compartment whereas contacts in the repressive B compartment decreased. Although still transcriptionally inactive, genes moving away from the nucleolus in NPCs were implicated in neuron development and differentiation processes, indicating that the detachment from the nucleolus marks a first step toward activation in later stages of differentiation. The results highlight the role of the nucleolus as repressive compartment that is implicated in the control of gene expression program during lineage commitment. Finally, the methodologies here developed for the identification of NADs at genome-wide level will now make possible to include the contribution of the nucleolus in future studies investigating the relationship between nuclear space and genome function.

## Results

### Establishment of Nucleolar-DamID

We thought to map NADs by establishing an alternative experimental procedure that is not based on biochemical isolation of nucleoli. We reasoned to adapt the DNA adenine methyltransferase identification (DamID) method, which was successfully used to identify LADs in many cell types (van Steensel et al., 2001). In this application, Lamin B1 was fused to Dam from *Escherichia coli*. When Lamin B1-Dam is expressed in cells, DNA in molecular contact with the NL is methylated at the N6 position of adenine (m6A) within GATC sequences and can be mapped. However, since the nucleolus is a membrane-free compartment, the application of DamID for the identification of NADs (Nucleolar-DamID) required further adaptations. A first criterion was to fuse Dam with proteins that are mainly localized within nucleolus and interact with DNA independently of the sequence content. We reasoned to exclude nucleolar proteins implicated in ribosome biogenesis (nucleolin, fibrillarin, UBF etc.) since their use might influence the readout of Nucleolar-DamID, such as adenine methylation of sequences only within the rRNA genes, thereby excluding the detection of other genomic domains located within nucleoli. Therefore, we thought to engineer a nucleolar histone, which can bind DNA sequences without motif specificity and has the ability to localize exclusively within nucleoli (**Fig. 1A**). To assess this possibility, we inserted a nucleolar localization signal (NoLS, RKKRKKK) (Birbach et al., 2004; Emmott and Hiscox, 2009) at the C-terminus of the histone H2B (H2B-NoLS). Live cell imaging of NIH3T3 cells transfected with a plasmid expressing GFP-tagged H2B-NoLS revealed a prominent and preferential localization in nucleoli when compared to the homogeneous nuclear distribution of H2B-GFP (**Fig. 1B**). Chromatin fractionation analyses showed that H2B-GFP-NoLS associated with chromatin similarly to H2B-GFP (**Fig. 1C**). Next, we tested whether the nucleolar localization of H2B-NoLS depends on the integrity of the nucleolus by inhibiting rRNA synthesis with Actinomycin D (ActD), a potent inhibitor of rRNA gene transcription. Downregulation of rRNA synthesis is known to induce a spatial reorganization of the nucleolar structure with the migration of its components, including rRNA genes, to the nucleolar periphery, forming the so called nucleolar caps (Reynolds et al., 1964). Treatment of NIH3T3 cells with ActD for 24 hours after transfection of plasmids expressing GFP-tagged proteins did not affect the localization of H2B-GFP whereas, as expected, the nucleolar transcription terminator factor I (TTF1), which associates with rRNA genes (Evers and Grummt, 1995), segregated into nucleolar caps (**Fig. 1D**). In contrast, upon ActD treatment, H2B-GFP-NoLS signal localized outside the nucleolus and appeared larger compared to the condensed structure of nucleolar caps (**Fig. 1D**). These results indicated that the nucleolar localization of H2B-NoLS depends on nucleolus integrity. Furthermore, the data suggest that genomic regions that incorporated H2B-NoLS in the nucleolus redistributed outside of it upon nucleolar segregation.

**Figure 1.**
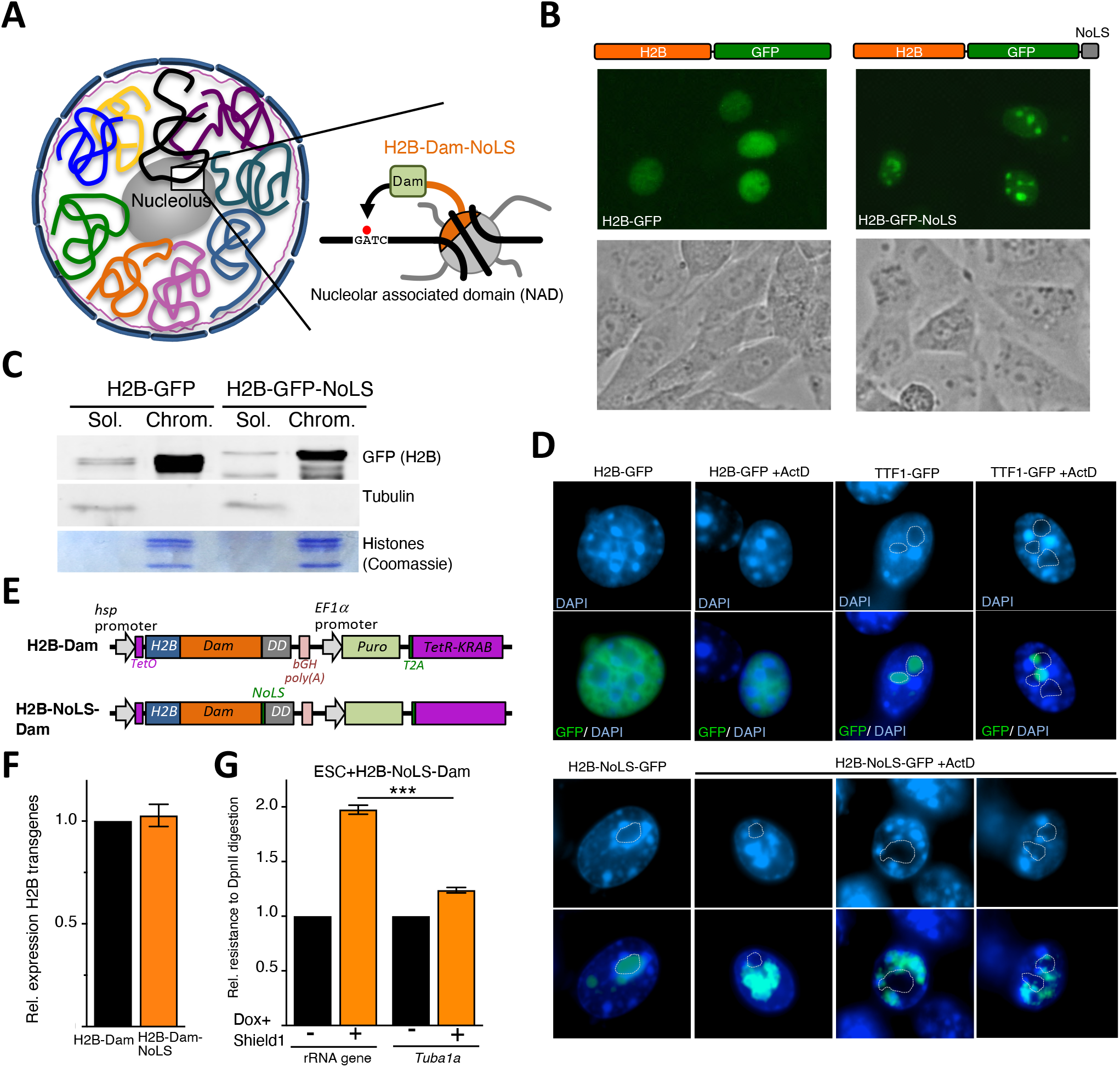
Establishment of Nucleolar-DamID. **A.** Scheme representing the strategy for the establishment of Nucleolar-DamID. **B.** Live cell imaging of NIH3T3 cells 24 hours post-transfection with plasmids expressing H2B-GFP and H2B-GFP-NoLS under the minimal CMV promoter. Phase contrast images serve to visualize the nucleoli. **C.** Chromatin-bound (Chrom.) and soluble (Sol.) fractions of equivalent number of NIH3T3 cells transfected with H2B-GFP and H2B-GFP-NoLS were analyzed by western blot using anti-GFP antibodies. Tubulin and histones are shown as loading and fractionation control. **D.** Representative immunofluorescence images of NIH3T3 cells transfected with H2B-GFP, TTF1-GFP, and H2B-NoLS-GFP and treated for 24 hours with or without ActD (50 ng/ml). ActD was added to cells 24 hours post-transfection. Nucleoli can be visualized by the low DAPI intensity and are highlighted with a dotted line. **E.** Scheme representing the construct used for double inducible expression of Dam-fused H2B and nucleolar H2B (H2B-NoLS) proteins. **F.** qRT-PCR showing similar expression levels of H2B-Dam and H2B-NoLS-Dam in ESCs. Data are from three independent experiments. Error bars represent s.d. **G.** m6A levels at rRNA genes and *Tuba1a* in ESCs with and without 15 hours treatment with 100 ng Doxycycline (Dox) and 1 μM Shield1. m6A levels were measured by digestion of genomic DNA with DpnII, which is blocked by m6A, followed by quantitative amplification with primers encompassing the Dam GATC element. Normalization was achieved through measurements with primers encompassing sequences lacking GATC. Data are from three independent experiments. Error bars represent s.d. Statistical significance (*P*-values) for the experiments was calculated using the paired two-tailed t-test (*** < 0.001).

To assess the efficacy of the H2B-NoLS in DamID application, we established mouse ESC lines with heterozygotic integration into the *Rosa26* locus of a transgene that allows inducible expression of H2B-Dam or H2B-Dam-NoLS chimeric proteins (**Fig. 1E, Fig. S1A**). In order to not saturate Dam sites, the expression of Dam-fused proteins was kept low by placing transgene transcription under the control of the minimal *hsp* promoter that was further modulated using a double inducible system for the control of transcription and protein stability. We used the TetR-KRAB repressor system by inserting two Tet operator (*TetO*) sequences downstream the minimal *hsp* promoter and an *EF1α* promoter*-Puro-T2A-TetR-Krab* cassette downstream the *Dam* transgenes. Furthermore, we placed at the C-terminus of both transgenes a destabilization domain (DD) that causes proteins to be rapidly targeted for proteasomal degradation unless the protein is shielded by the synthetic small molecule Shield1 (Banaszynski et al., 2006). The lack of leakiness of this inducible system was confirmed by the transfection of HEK 293T cells with plasmids containing the sequences *hsp-TetO-GFP-Dam-DD-EF1a-puro-T2A-TetR-KRAB* (**Fig. S1B**). Expression of H2B-Dam-DD and H2B-Dam-NoLS-DD transgenes in the corresponding ESC lines was induced for 15 hours of treatment with doxycycline (Dox) and Shield1. The low expression of both transgenes did not allow measurements of protein levels by western blot but only quantifications by qRT-PCR, which showed that H2B-Dam-DD and H2B-Dam-NoLS-DD were expressed at similar levels (**Fig. 1F**). To test the specificity of H2B-Dam-NoLS in depositing m6A at nucleolar sequences, we measured GATC methylation levels at rRNA genes, which are located in nucleoli, and *Tuba1a*, which we did not expect to be located in nucleoli (**Fig. 1G**). We found significantly higher m6A levels at rRNA genes compared to *Tuba1a*, indicating that the nucleolar H2B histone was preferentially incorporated in rRNA genes, the known genetic component of the nucleolus.

### Nucleolar-DamID identifies NADs

The identification of LADs by DamID was obtained by measuring the m6A ratio of Lamin B1-Dam over a freely diffusible Dam fused to GFP (GFP-Dam) (Peric-Hupkes et al., 2010). However, in the Nucleolar-DamID, we opted to use as a control the measurement of m6A levels in ESCs expressing H2B-Dam. This strategy was based on the fact that compared to GFP-Dam, which has also been reported to have some preference for open chromatin (Aughey et al., 2018), the two Dam-fused histones should only differ in their nuclear localization. Furthermore, the use of H2B-Dam as control will also serve to compensate for the eventual incorporation of nucleolar histones in genomic regions outside the nucleolus.

We combined two independent Nucleolar-DamID experiments, which were highly correlated (Pearson correlation 0.89) (**Fig. S2A**). We used a previously established DamIDseq pipeline (Marshall and Brand, 2015) and constructed genome-wide maps of NADs in ESCs by taking the resulting H2B-Dam-NoLS over H2B-Dam m6A ratio as a measure for the relative contact frequency of DNA sequences with the nucleolus (FDR <0.01) (**Fig. 2A, Fig. S2B**). We found that nucleolar H2B contacts in ESCs display broad domains ranging between 70 kb and 6.2 Mb (**Fig. S2C**). These contacts (from here on termed as NADs) were distributed all over the chromosomes with some preferences for chromosomes containing rRNA genes (**Fig. 2B,C, Fig. S2B**). The X chromosome, which is active in the male ESCs used in this study, showed the lowest enrichment in NADs. Consistent with the well-known co-localization of centromeres in the vicinity of the nucleolus (Padeken and Heun, 2014), we found that NADs were enriched in the centromere-proximal regions of the large majority of chromosomes, which in the mouse cells are all acrocentric (**Fig. 2B**). These data are also consistent with recent results based on the SPRITE method that identified a sub-class of NADs interacting with rRNA transcripts, which are synthesized in the nucleolus, and are in close linear proximity to centromeres (Quinodoz et al., 2018). Furthermore, the Nucleolar-DamID identified as NADs all sequences that SPRITE found to form inter-chromosomal contacts that interact with rRNA (i.e. nucleolar hub) (Quinodoz et al., 2018) (**Fig. S2C**). To further support the specificity of the Nucleolar-DamID, we recovered reads containing rRNA gene (rDNA) contacts from a high resolution (<750 bp) Hi-C map in ESCs, the highest to date in mammalian cells (Bonev et al., 2017) (from here on named as HiC-rDNA). The adaptation of HiC analysis for the identification of rDNA contacts was based on the modification of the mouse reference genome, which does not contain rRNA gene sequences, by adding at the end of chromosome 12 one copy of rRNA gene unit (see methods). As in the case of NADs identified by SPRITE (Quinodoz et al., 2018), rDNA contacts obtained with HiC-rDNA should represent a sub-class of NADs since not all genomic domains associating with the nucleolus must necessarily interact with the rRNA genes. The most frequent rDNA contacts were found at chromosomes containing rRNA genes (**Fig. 2D,E**, **Fig. S3A**), an expected result since these interactions mainly occur in *cis*. Furthermore, rDNA contacts were enriched at centromeric-proximal regions of all chromosomes (**Fig. 2E,F**, **Fig. S3A**), suggesting that the known co-localization of centromeres in the vicinity of the nucleolus might promote the formation of rDNA interactions between different chromosomes. Importantly, 76% of rDNA contacts identified by HiC-rDNA were located in NADs found with the Nucleolar-DamID (**Fig. 2G,H**). Thus, the Nucleolar-DamID can identify a large portion of rDNA contacts, a sub-class of NADs identified by HiC-rDNA. Finally, we measured the cellular localization of a selected NAD on chromosome 19 by FISH and found that all analyzed cells display a signal in close proximity or within the nucleolus (**Fig. 2I**). In contrast, a FISH probe hybridizing against a DNA region corresponding to LAD and not mapped as NAD was mainly localized at the NL (**Fig. S2D**). Together, the data showed that Nucleolar-DamID is an accurate method for genome-wide identification of all NADs.

**Figure 2.**
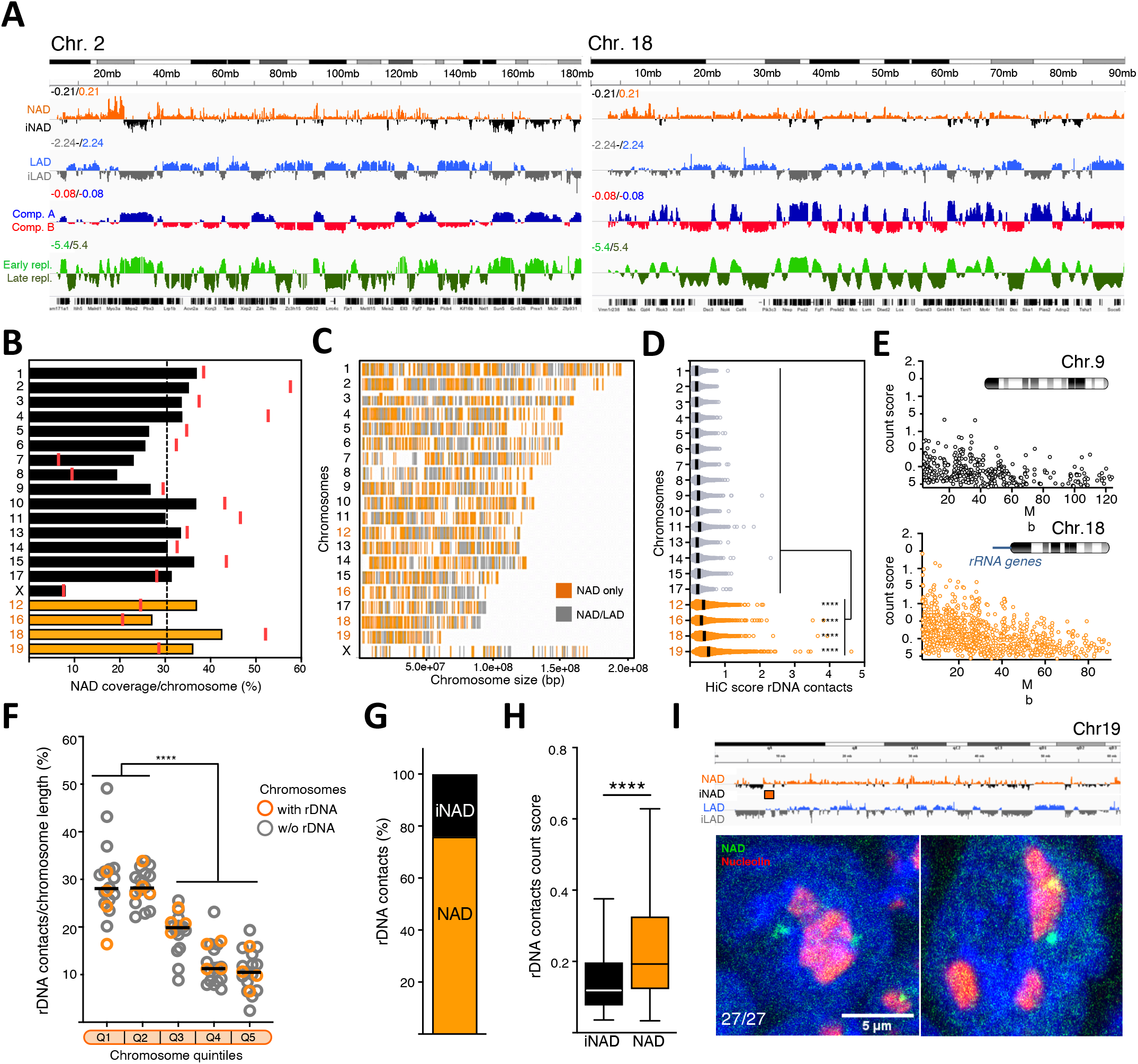
Nucleolar-DamID identifies NADs. **A.** Chromosomal view of NADs, LADs (Peric-Hupkes et al., 2010), A and B compartments (Dalcher et al., 2020), and early and late replicating DNA (Marchal et al., 2018) in ESCs. NADs are measured as log_2_ ratio of m6A levels between H2B-Dam-NoLS and H2B-Dam. iNAD: regions not contacting the nucleolus. iLAD: regions not contacting the NL. Chromosomes 2 and 18 are shown. **B.** NAD coverage. Bars represent NAD coverage values of each mouse chromosome. Red lines showed NAD coverage values in the centromere-proximal end of each chromosome. Chromosome bearing rRNA genes are depicted with orange numbers and bars. Dotted line shows NAD whole genome coverage. **C.** Nucleolar H2B-Dam chromosomal interaction maps. NAD-only and NAD/LAD regions were represented with orange and grey colors, respectively. Chromosome bearing rRNA genes are depicted with orange numbers. **D.** Genomic contacts with rRNA genes (rDNA contacts) obtained from published HiC maps (Bonev et al., 2017). Data represent the proportion of identified unique contacts for each chromosome. **E.** Representative images showing normalized count score of rDNA contacts obtained from the HiC-rDNA on chromosome 9, which does not harbor rRNA genes, and chromosome 18, which contains rRNA genes. **F.** Distribution of rDNA contacts for each chromosome. Values represent the proportion of the number of unique contacts for each chromosome quintile. Statistical significance (*P*-values) for the experiments was calculated using the unpaired two-tailed t-test (**** < 0.0001). **G.** NADs identified by Nucleolar-DamID are enriched in rDNA contacts. Data represent the proportion of identified unique HiC-rDNA contacts at NADs and regions non contacting the nucleolus (iNAD). **H.** Hi-C normalized count score of identified unique rDNA contacts at NADs and iNADs. Statistical significance (*P*-values) for the experiments was calculated using the unpaired two-tailed t-test (**** < 0.0001). **I.** Upper panel. NAD and LAD composition of chromosome 19 and the NAD FISH probe (orange bar). Lower panel. Example images from immunofluorescence for nucleolin (red) combined with DNA FISH using probes corresponding to NAD-only region of chromosome 19 (green), and DAPI (blue). Numbers refer to cells showing the signal of NAD probe located close to or within the nucleoli relative to the number of all analyzed ESCs (27/27).

### Distinct layers of repressive chromatin states distinguish genomic domains according to their interaction with the nucleolus, nuclear lamina, or both

To gain insights on how genome compartmentalization at nucleolus and NL is related to each other, we analyzed the NAD composition relative to LADs, which were previously identified in ESCs using LaminB1-DamID (Peric-Hupkes et al., 2010). We found that about 53% of NADs correspond to LADs whereas 40% of LADs are also NADs (**Fig. 3A,B**). We termed this NAD subclass NAD/LAD whereas NADs not overlapping with LADs were named NAD-only. Accordingly, the average Lamin B1-Dam signal highly increased at NAD/LAD boundaries and to a much less degree with NAD-only boundaries (**Fig. 3C**). These results support previous data showing that some LADs can also be found in the vicinity of the nucleolus (Kind et al., 2013; Ragoczy et al., 2014). NAD-only, NAD/LAD, and LADs that do not overlap with NAD (LAD-only) have low gene density (**Fig. 3D**) and, consistent with previous reports, NADs are particularly enriched in olfactory receptor genes and zinc finger genes (Nemeth et al., 2010; van Koningsbruggen et al., 2010). Both NAD subclasses and LAD-only showed distinct chromatin and transcriptional features. A large portion of NAD-only regions localized in the active A compartment (66%), was early replicating (65%), and had higher gene density relative to NAD/LAD and LAD-only sequences (**Fig. 3D-F**). However, NAD-only had lower gene density relative to the whole genome and low level of gene expression compared to genes located in the active A compartment, indicating that localization at the nucleolus correlates with low gene activity (**Fig. 3D,G**). In contrast to NAD-only, the majority of NAD/LAD and LAD-only were located in the repressive B compartment (90% and 80%) and were late replicating (82% and 63%), a result that is consistent with previous data showing that total LADs are characterized by these repressive features (Kind et al., 2015; Peric-Hupkes et al., 2010; Pope et al., 2014) (**Fig. 3D-F**). However, compared to LAD-only, NAD/LAD appeared to be more enriched in the B compartment and in late-replicating DNA regions and displayed lower gene density and gene expression levels (**Fig. 3D-G**), suggesting that sequences that can localize at both nucleolus and NL have enhanced repressive chromatin features than sequences that anchor only to NL. These results were also supported by the different levels of active and repressive histone marks at these genomic domains (**Fig. 3H**). As expected, NAD-only sequences had higher levels of the active histone modifications H3K4me3, H3K27ac, and H3K4me1 compared to LAD-only and NAD/LAD. However, and consistent with the low gene expression (**Fig. 3G**), NAD-only contain lower levels of active histone marks compared to genomic regions within the A active compartment.

**Figure 3.**
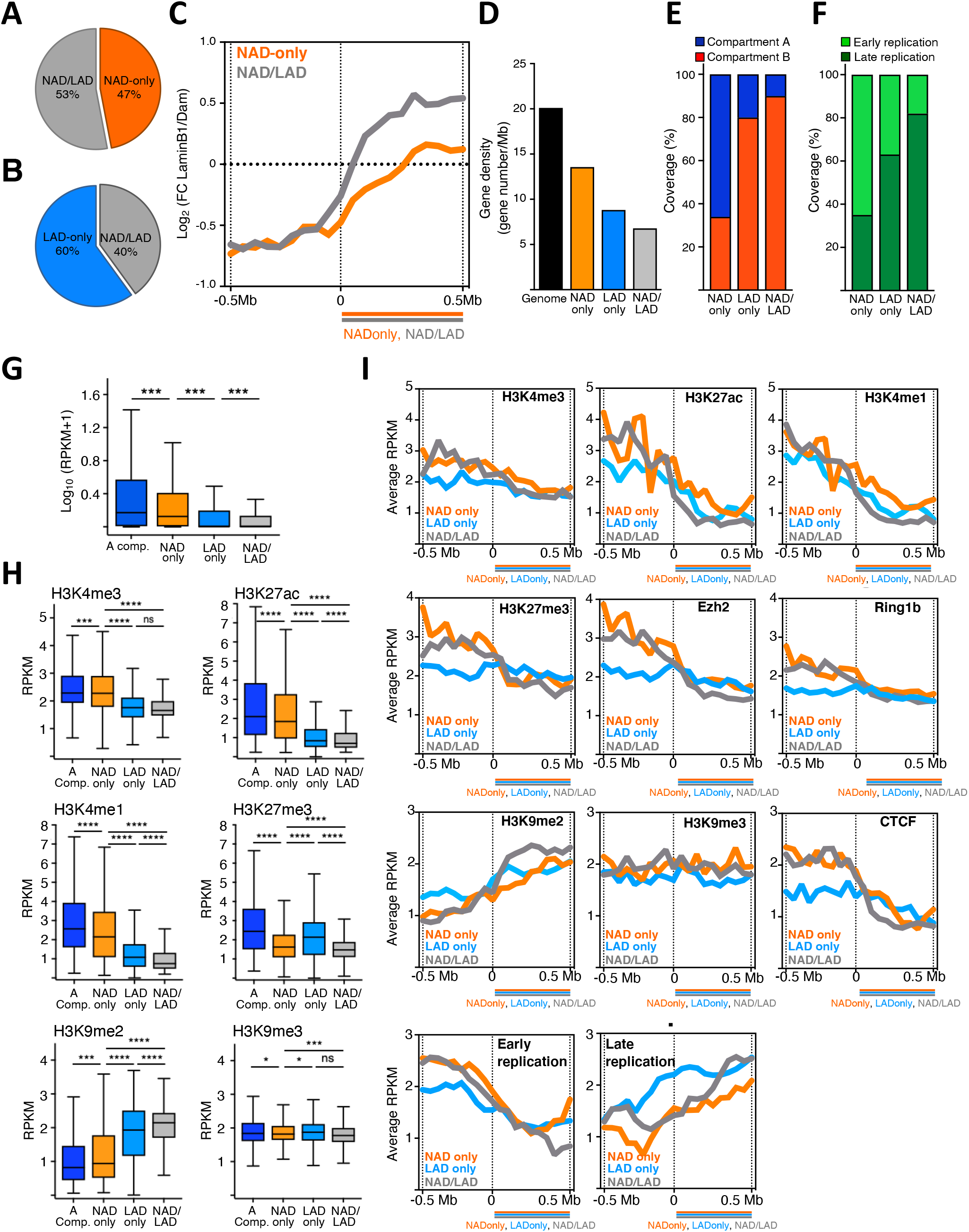
Distinct layers of repressive chromatin states distinguish genomic domains according to their interaction with the nucleolus, nuclear lamina, or both. **A.** Venn diagram showing the proportion of NAD-only and NAD/LAD regions in NADs identified by Nucleolar-DamID. **B.** Venn diagram showing the proportion of LAD-only and NAD/LAD regions in LADs of ESCs. **C.** Lamin B1-DamID scores plotted over NAD-only and NAD/LAD boundaries in ESCs. **D.** Gene density of total genome, NAD subclasses, and LAD-only. **E.** Amounts (%) of NAD-only, LAD-only and NAD/LAD in A and B compartments. **F.** Amounts (%) of early and late replicating DNA of NAD-only, LAD-only and NAD/LAD sequences. **G.** Expression values (RPKM) of genes within A compartment (A Comp.), NAD-only, LAD-only, and NAD/LAD. Statistical significance (*P*-values) was calculated using the unpaired two-tailed t-test (***< 0.001). **H.** Levels of active histone marks (H3K4me3, H3K27ac, H3K4me1) and repressive histone marks (H3K27me3, H3K9me2, H3K9me3) at genomic regions located at the A compartment (A Comp.) and NAD-only, LAD-only, and NAD/LAD regions. Dataset used in this analysis are listed in **Table S12**. Values are shown as average RPKM. Statistical significance (*P*-values) was calculated using the unpaired two-tailed t-test (*<0.05, ***< 0.001, ****< 0.0001, ns: non-significant). **I.** Occupancy (average RPKM) of histone modifications, Ezh2, Ring1b, CTCF and early and late-DNA replication plotted over the boundaries of NAD-only (orange lane), NAD/LAD (blue line), and NAD/LAD (grey line), respectively.

Furthermore, relative to A compartment, NAD-only were depleted of H3K27me3 and enriched in H3K9me2 (**Fig. 3H**), suggesting that the localization close to the nucleolus marks transcriptionally repressive states that might not depend on Polycomb. We also observed that NAD/LAD and LAD-only regions showed distinct chromatin features. NAD/LAD displayed a more repressive chromatin state than LAD-only. They contained a lower amount of active histone marks and, in particular, higher levels of H3K9me2 whereas H3K9me3 were similar (**Fig. 3H**). Furthermore, the levels of the facultative heterochromatin mark H3K27me3 were lower in NAD/LAD than in LAD-only, indicating that contacts with the nucleolus, even for sequences also able to interact with the NL, shape a specific repressive chromatin state that is characterized by enrichment in H3K9me2 and low H3K27me3 content. These results indicate that genomic regions display a distinct repressive chromatin composition according to their ability to localize at the nucleolus, NL, or both. As in the case of LADs, both NAD-only and NAD/LAD were characterized by abrupt borders, which display a sharp transition in several chromatin features (**Fig. 3I**). The average signal of active histone marks, CTCF and early replicating DNA sharply decreased at the border of both NAD subclasses and LAD-only. These results further indicate that contacts with the nucleolus demarcate less active chromatin domains. In contrast to LADs, which were described to be enriched in H3K27me3 and Polycomb components Ezh2 and Ring1b near LAD borders (Harr et al., 2015), NAD-only and NAD/LAD did not show this enrichment but instead they displayed a more drastic decrease of these factors at their corresponding borders (**Fig. 3I**). H3K9me2 levels and late DNA replication signals sharply increased at the borders of both NAD sub-classes and LAD-only whereas H3K9me3 did not show this trend. Thus, the identification of NADs with the Nucleolar-DamID allowed to distinguish different layers of genome compartmentalization by defining regions that are exclusively localized at nucleoli, NL, or both. Further, the results revealed that NADs demarcate regions of the genome with a repressive state and specifically enriched in H3K9me2, linking this modification with nuclear architecture.

### The chromatin state of rRNA genes regulates H3K9me2 levels at sequences adjacent to the nucleolus

Previous work showed that the chromatin state of rRNA genes could affect chromatin structures outside the nucleolus (Savić et al., 2014). In ESCs, all rRNA genes are euchromatic due to the impairment of processing of the long non-coding IGS-rRNA into mature pRNA, which is required for the recruitment of the repressive nucleolar remodeling complex NoRC to rRNA genes (Dalcher et al., 2020; Guetg et al., 2012; Mayer et al., 2006; Savić et al., 2014). The active state of rRNA genes in ESCs could be reversed by addition of mature pRNA that caused NoRC recruitment and consequent formation of heterochromatin at rRNA genes, including the increase in H3K9me2 and H3K9me3 (**Fig. 4A,B**) (Leone et al., 2017; Savic et al., 2014). This heterochromatinization was not only limited to rRNA genes but also it extended to other regions of the genome outside the nucleolus, such as minor satellites, highlighting a crosstalk between nucleolar and nuclear chromatin (**Fig. 4B**). The identification of NADs in ESCs prompted us to identify which genomic regions were affected by the chromatin state of rRNA genes and their relationship with the nucleolus. We measured H3K9me2 and H3K9me3 by ChIPseq of ESCs transfected with mature pRNA (**Fig. 4C, Fig. S4A**). We found that H3K9me2 levels increased at several genomic regions all over the chromosomes of ESC+pRNA compared to control cells whereas H3K9me3 levels were not affected (**Fig. 4C,D**). We observed a sharp increase in H3K9me2 at regions adjacent to the border of NADs but not of LADs, indicating an expansion of H3K9me2 domains specifically at regions neighboring NADs (**Fig. 4E,F**). These results revealed that the nucleolus is not only a scaffold where repressive chromatin can be positioned but also it is part of the regulatory process for the establishment or maintenance of repressive chromatin states.

**Figure 4.**
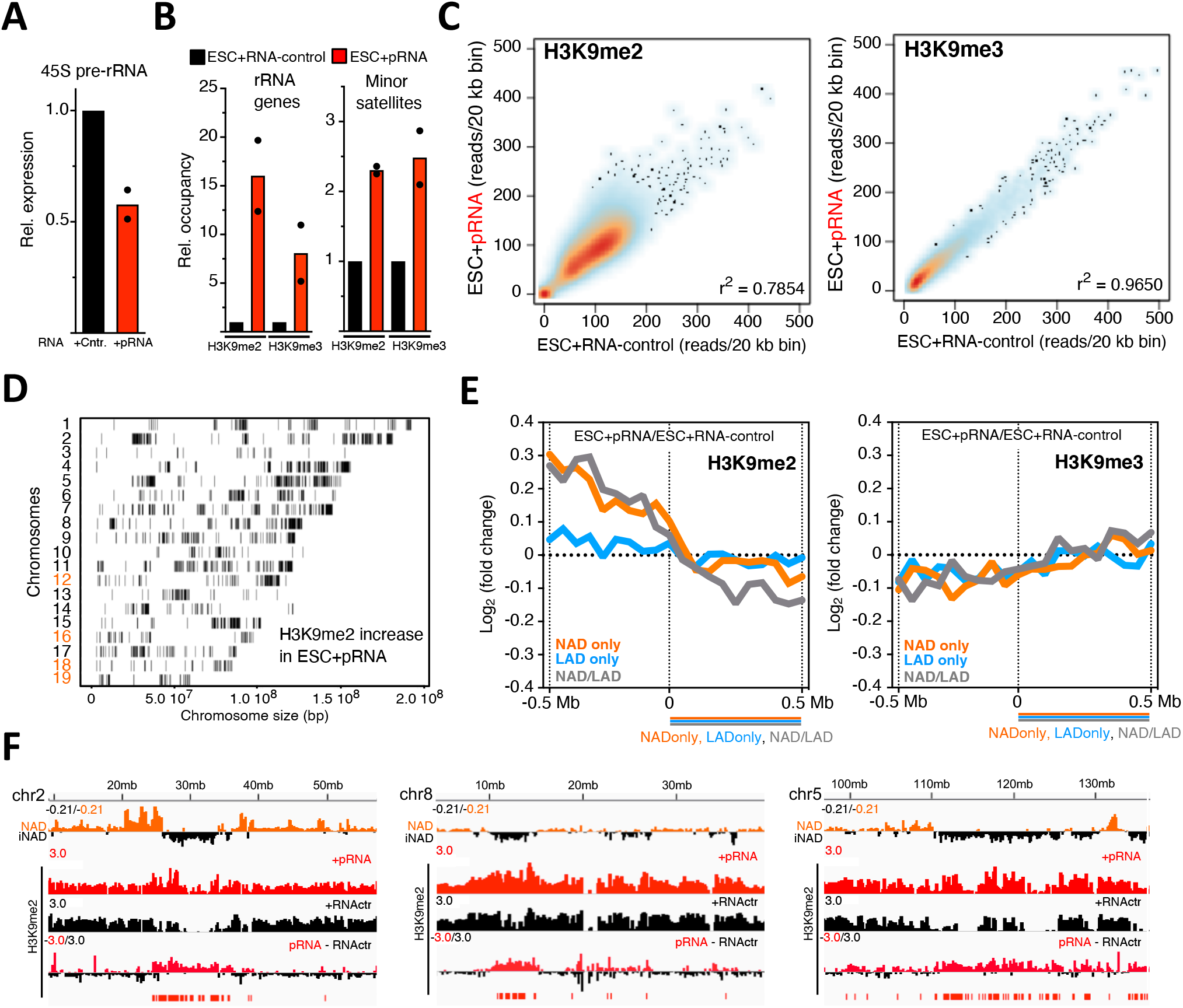
The chromatin state of rRNA genes regulates H3K9me2 levels at sequences adjacent to the nucleolus. **A.** qRT-PCR showing 45S pre-rRNA levels in ESCs transfected with pRNA or RNA-control. Values were normalized to *β-actin* mRNA and to ESCs transfected with RNA-control. Data are from two independent experiments. **B.** H3K9me2 and H3K9me3 ChIP in ESCs transfected with pRNA or RNA-control. Data were measured by qPCR and normalized to input and ESC+RNA-control. Data are from two independent experiments. **C.** Addition of pRNA in ESCs caused an increase in H3K9me2 at several genomic regions. Scatter plot showing H3K9me2 and H3K9me3 levels (reads/20kb bin) between ESC+pRNA and ESC+RNA-control. **D.** Chromosomal interaction map showing the distribution of regions with increased H3K9me2 levels in ESC+pRNA compared to ESC+RNA-control. **E.** Heterochromatinization of rRNA genes promotes H3K9me2 expansion at regions neighboring NADs. H3K9me2 fold changes in ESC+pRNA vs. ESC+RNA-control plotted over the boundaries of NAD-only (orange), LAD-only (blue), and NAD/LAD (grey). **F.** Representative images showing the increase of H3K9me2 at regions neighbouring NADs.

### Dynamics of genome organization around the nucleolus during early development

Next, we asked whether NAD composition could change according to developmental stage. We performed Nucleolar-DamID and HiC-rDNA analyses in neural progenitor cells (NPCs) derived upon differentiation of ESCs using established protocols (Bibel et al., 2004). Both Nucleolar-DamID and HiC-rDNA revealed a common, global chromosome architecture between ESCs and NPCs, with substantial overlapping interactions with the nucleolus (**Fig. 5A,B, Fig. S2A**). However, there were remarkable differences in the organization of the chromosomes around the nucleolus of ESCs and NPCs. Both Nucleolar-DamID and HiC-rDNA revealed that in all chromosomes contacts with the nucleolus were more frequent in NPCs than in ESCs (**Fig. 5C,D, Fig. S3A, S5A**). However, NAD coverage and the number of unique rDNA contacts was higher in ESCs than in NPCs (**Fig. 5E, Fig. S5B**). Furthermore, a large fraction of unique rDNA contacts in NPCs were located at chromosomes bearing rRNA genes whereas in ESCs the distribution of contacts was more homogeneous among the chromosomes compared to NPCs. Finally, rDNA contacts were more enriched at the centromere-proximal regions of all chromosomes in NPCs than in ESCs (**Fig. 2F, 5F**). These results suggest that the architecture of chromosomes surrounding the nucleolus in NPCs is in a more compact, rigid form, making fewer but more frequent contacts with rRNA genes. In contrast, the structure of chromosomes around the nucleolus of ESCs appears more flexible, establishing more but less frequent rDNA-contacts. Next, we investigated for the presence of genomic contacts with the nucleolus that are specific for either ESCs or NPCs. We identified large domains that contained rDNA contacts exclusively in ESCs or NPCs (here after referred as ESC_sp_- or NPC_sp_-rDNA contacts; average length 0.51 Mb and 0.23 Mb, respectively; **Fig. 5G**). The large majority (>80%) of ESC_sp_- and NPC_sp_-rDNA contacts were located in the repressive B compartment, a result that further links the nucleolus with repressive chromosome structure (**Fig. 5H**). Analysis of eingenvector values revealed that ESC_sp_-rDNA contacts increased in NPCs the interaction strength within the active A compartment whereas contacts in the repressive B compartment decreased (**Fig. 5G,I,J**). 37% of ESC_sp_-rDNA contacts switched from B to A compartment in NPCs and ESC_sp_-rDNA contacts that remained in B (47%) or A compartments (16%) in NPCs significantly increased their eigenvector values in NPC (**Fig. 5I**). Similar results were observed for NPC_sp_-rDNA contacts in ESCs. Thus, rDNA contacts specific to ESCs or NPCs correspond to cell-type specific repressive features of the genome organization. To determine whether changes in rDNA contacts between undifferentiated and differentiated cells correspond to gene expression changes, we performed RNAseq of ESCs and NPCs. The majority of genes located at ESC_sp_- and NPC_sp_-rDNA contacts were lowly expressed (< 1 RPKM) and we did not observe significant changes in gene expression between ESCs and NPCs (**Fig. 5K**), indicating that the detachment from the nucleolus is not sufficient to reactivate gene expression. However, we found that the top 10 gene ontology (GO) terms for genes located in ESC_sp_-rDNA contacts were linked to pathways implicated in neuron development and differentiation (**Fig. 5L**), suggesting that the relocation of these genes away from the nucleolus might be a first step toward their activation in later stages of differentiation. On the other hand, genes located at NPC_sp_-rDNA contacts were linked to pathways of sensory perception of smell (mainly due to the presence of several olfactory receptor genes) and metabolic processes.

**Figure 5.**
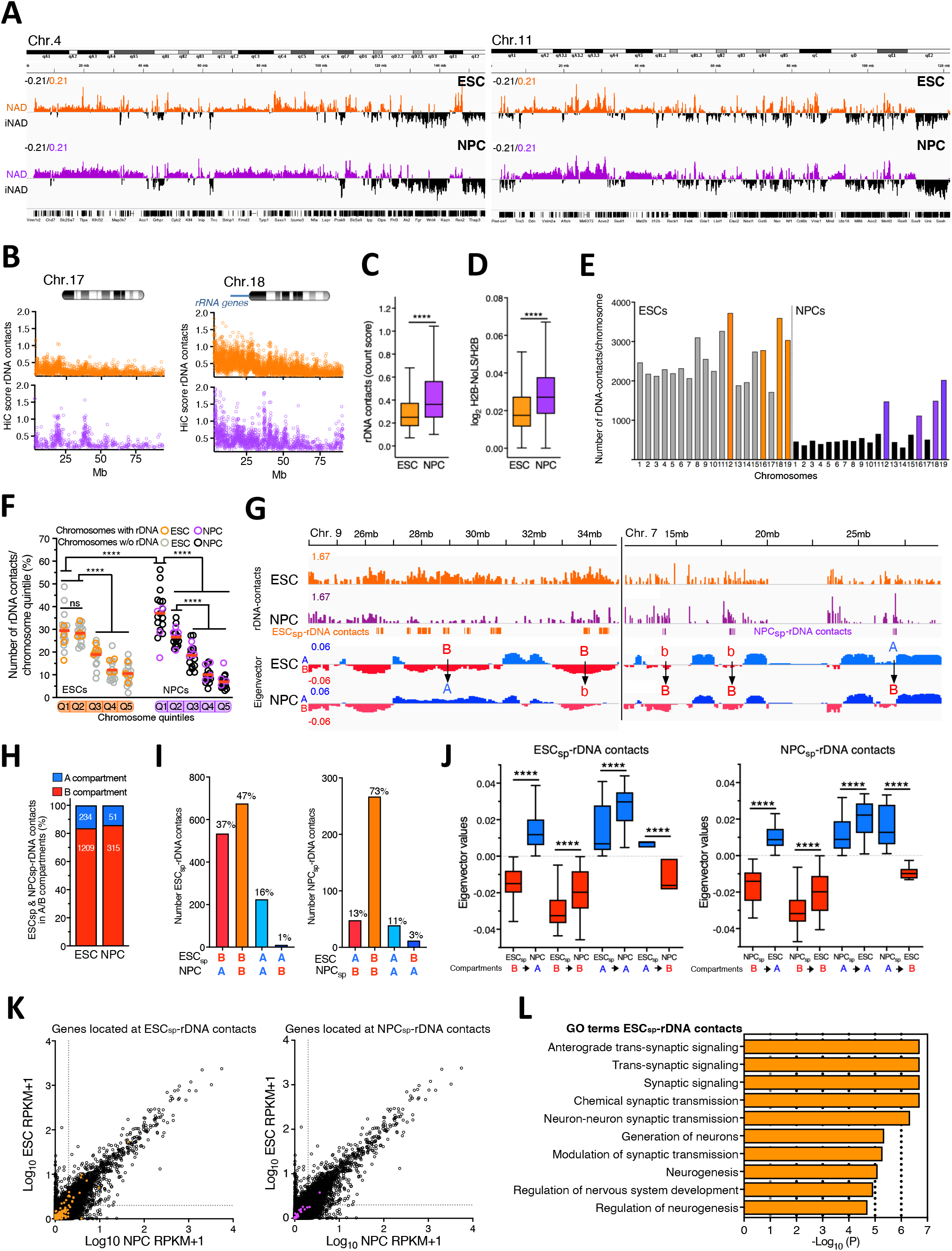
ESC and NPC differ in their chromosome organization around the nucleolus. **A.** Nucleolar-DamID. Chromosomal view of NADs in ESCs and NPCs. NADs are measured as log_2_ ratio of m6A levels between H2B-Dam-NoLS and H2B-Dam. iNAD: regions not contacting the nucleolus. iLAD: regions not contacting the NL. Chromosome 4 and 11 are shown. **B.** HiC-rRNA. Representative images showing HiC-score of rDNA contacts at chromosome 7, which does not harbor rRNA genes, and chromosome 18, which contains rRNA genes close to centromere (see also **Fig. S3**). **C,D.** Genomic contacts with the nucleolus are more frequent in NPCs than in ESCs. HiC-score of rDNA contacts (**C**) and Nucleolar-DamID values of NADs (**D**) in ESCs and NPCs. Statistical significance (*P*-values) was calculated using the unpaired two-tailed t-test (****< 0.0001). **E.** ESCs have more rDNA contacts than NPC. Number of unique rDNA contacts at each chromosome in ESCs and NPCs. Orange and violet bars refer to chromosomes containing rRNA genes in ESCs and NPCs, respectively. **F.** rDNA contacts are enriched in the centromeric-proximal regions of all chromosomes of NPCs relative to ESCs. To allow a better comparison, the data of ESCs in **Figure 2F** were plotted together with the data of NPCs. Values represent the proportion of rDNA-contacts for each chromosome quintile of ESCs and NPCs. Statistical significance (*P*-values) was calculated using the unpaired two-tailed t-test (****< 0.0001). **G.** Representative images of ESC_sp_- and NPC_sp_-rDNA contacts and their eigenvector values for A and B compartment. Arrows highlight changes in eigenvector values. B to A and A to B represent switch of compartments. B to b and b to B indicates a decrease or increase of eigenvector values between ESCs and NPCs. **H.** ESC_sp_- and NPC_sp_-rDNA contacts are in the repressive B compartment. Amounts (%) of ESC_sp_- and NPC_sp_-rDNA contacts in A and B compartments. **I.** Values represent the number of ESC_sp_- and NPC_sp_-rDNA contacts and their corresponding location in A and B compartments of ESCs and NPCs. **J.** Box plots showing eigenvector values of ESC_sp_- and NPC_sp_-rDNA contacts in the active A (blue) and repressive B (red) compartments. Statistical significance (*P*-values) was calculated using the paired two-tailed t-test (****< 0.0001). **K.** Scatter plot showing expression levels between ESC and NPCs. Expression of genes located at ESC_sp_- and NPC_sp_-rDNA contacts are highlighted in orange and magenta, respectively, whereas total genes are represented in black. Dotted lines indicate RPKM value as 1. **L.** Top 10 gene ontology terms of genes located at ESC_sp_-rDNA contacts.

Since rDNA contacts represent a sub-group of NADs, we extended our analyses to all NADs defined by Nucleolar-DamID. We identified NADs specific for ESCs or NPCs (here after defined as ESC_sp_-NAD and NPC_sp_-NAD). We found that ESCs have more cell type specific NADs than NPCs (16% and 8% coverage relative to the corresponding total NADs), underscoring a differential spatial organization of chromosomes around the nucleolus between the two cell types. The large majority of ESC_sp_-NAD-only (78%) lost in NPCs their contacts with both nucleolus and NL (iNAD/iLAD) whereas 66% of ESC_sp_-NAD/LAD became exclusively localized at NL (LAD-only) (**Fig. 6A**). Similar changes were observed with NPC_sp_-NAD in ESCs. We also analyzed whether cell types specific LAD-only could change their cellular location relative to the nucleolus and found that 91% of ESC_sp_-LAD-only in NPCs and 81% of NPC_sp_-LAD-only in ESCs lost their contacts with NL and did not associate with the nucleolus (**Fig. 6B**). These results suggest that the location of cell-type specific NADs and LADs at nucleolus, NL, or both influences the cellular location in the other cell type. Genomic regions exclusively contacting the nucleolus or the NL tend to lose contacts with both repressive compartments. In contrast, cell-type specific genomic regions that contact both nucleolus and NL (NAD/LAD) preferentially switch their cellular position toward the nuclear periphery.

**Figure 6.**
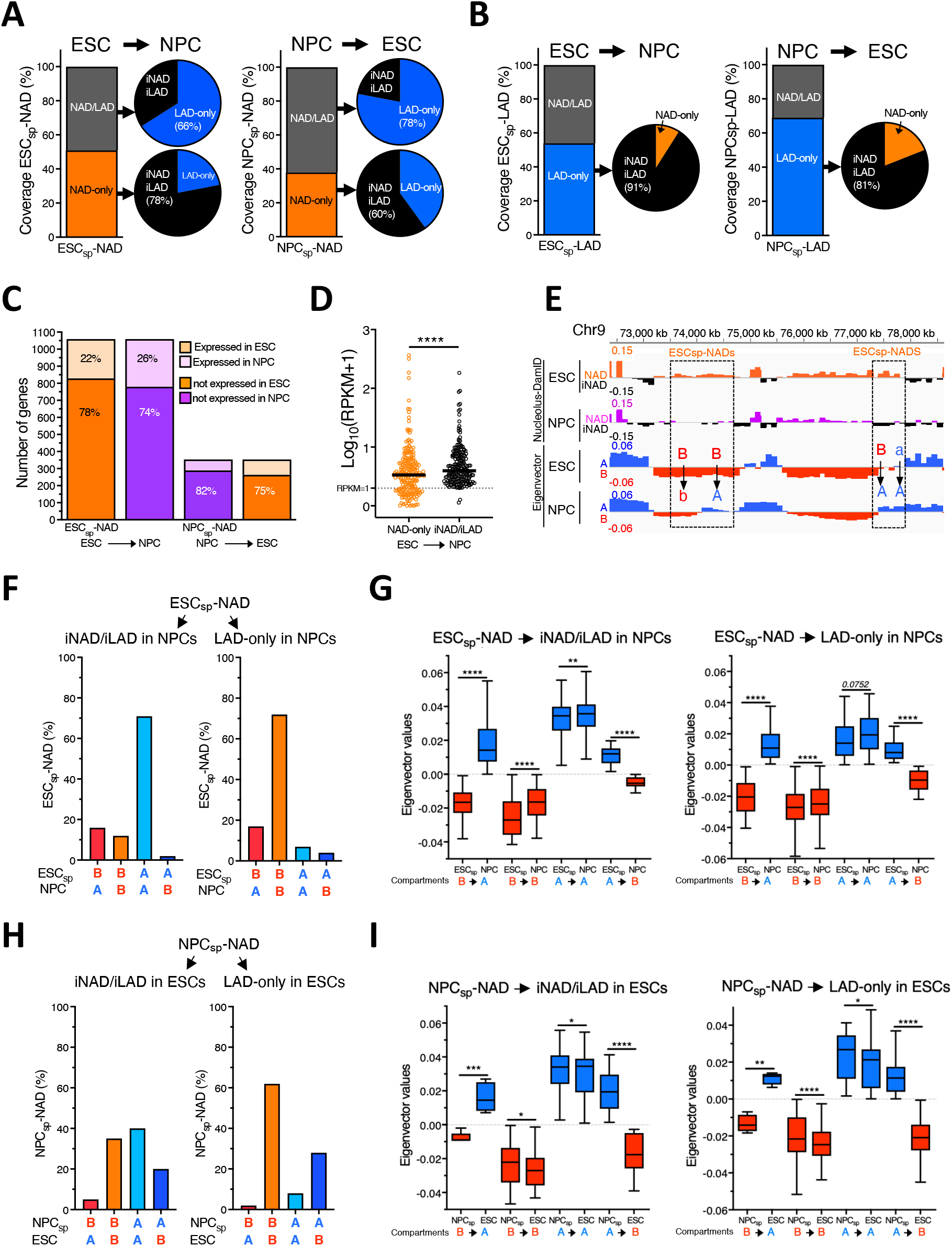
NAD organization in ESCs and NPCs. **A.** Coverage of ESC_sp_- and NPC_sp_-NAD types (NAD/LAD and NAD-only) in ESCs and NPCs. The location of ESC_sp_- and NPC_sp_-NAD types as iNAD/iLAD and LAD-only in NPCs and ESCs, respectively, and the proportion are represented with Venn diagrams. **B.** Coverage of ESC_sp_- and NPC_sp_-LAD types (NAD/LAD and LAD-only) in ESCs and NPCs. The location of ESC_sp_- and NPC_sp_-LAD types as iNAD/iLAD and NAD-only in NPCs and ESCs, respectively, and the proportion are represented with Venn diagrams. **C.** Number of genes located at ESC_sp_- and NPC_sp_-NAD. The proportion of low or not expressing (<1 RPKM) and expressing genes in ESCs and NPCs is indicated. **D.** Expression levels of genes located at ESC_sp_-NAD-only that lost contact with the nucleolus and nuclear lamina (iNAD/iLAD) in NPCs. Values are from genes that were expressed (> RPKM 1) in ESCs or NPCs. Statistical significance (*P*-values) was calculated using the paired two-tailed t-test (****< 0.0001). **E.** Representative images of ESC_sp_-NAD and their eigenvector values for A and B compartment. Arrows highlight changes in eigenvector values between ESCs and NPCs. B to A represents the regions switching from B (ESCs) to A (NPCs) compartment. B to b and a to A indicate higher eigenvector values in ESCs compared to NPCs. **F.** Values represent the number of ESC_sp_-NAD and their corresponding location in A and B compartments of ESCs and NPCs classified according to regions that move away from the nucleolus and nuclear lamina (iNAD/iLAD) or gain exclusive location at the nuclear periphery (LAD-only) in NPCs. **G.** Box plots showing eigenvectors values of ESC_sp_-NADs in the active A (blue) and repressive B (red) compartments. Statistical significance (*P*-values) was calculated using the paired two-tailed t-test (**\*\***< 0.01, ****< 0.0001). **H.** Values represent the number of NPC_sp_-NAD and their corresponding location in A and B compartments of ESCs and NPCs classified according to regions that move away from the nucleolus and nuclear lamina (iNAD/iLAD) or gain exclusive location at the nuclear periphery (LAD-only) in ESCs. **I.** Box plots showing eigenvectors values of NPC_sp_-NADs in the active A (blue) and repressive B (red) compartments. Statistical significance (*P*-values) was calculated using the paired two-tailed t-test (*< 0.05, **<0.01, **\*\*\***< 0.001, ****< 0.0001).

Genes located at ESC_sp_- and NPC_sp_-NADs (1201 and 377 genes, respectively) were either not or low expressing in both cell types (74-82% of genes with RPKM <1) and showed in general no significant changes in gene expression between ESCs and NPCs (**Fig. 6C, Fig. S6A**). The only exception was for genes located in ESC_sp_-NAD-only that became iNAD/iLAD in NPCs. A large fraction of these genes (221 out of 500 genes) were expressed in ESCs or in NPCs and displayed a modest but significantly lower expression in ESCs relative to NPCs (**Fig. 6D**). Remarkably, independently of gene expression states, these genes showed significant enrichment in many pathways linked to neuron development and differentiation (**Fig. S6B**), suggesting that their detachment from both repressive nucleolar and NL compartments might unlock them for activation in a next differentiation step. Accordingly, genes in ESC_sp_-NAD that repositioned exclusively at NL (LAD-only) in NPCs were mainly enriched in pathways linked to keratinocyte and epidermal differentiation (**Fig. S6C**), which should not be activated later in neurogenesis and thus remained in contact with a repressive compartment such as NL. These results further support a role of the nucleolus as repressive compartment that is implicated in the control of gene expression program during lineage commitment.

Consistent with the analysis of rDNA contacts, ESC_sp_-NAD in NPCs increased the interaction strength within the A compartment whereas contacts in the B compartment decreased (**Fig. 6E-G**). In contrast, we found that eigenvector values of NPC_sp_-NAD were higher in NPCs than in ESCs (**Fig. 6H,I**), suggesting that NPC_sp_-NAD represent a sub-type of chromatin organization that showed less interaction with the repressive B compartment relative to ESCs while keeping genes in an inactive expression state in both cell types.

Together, the results revealed the contribution of the nucleolus in the genome compartmentalization and function during early development.

## Discussion

The nucleolus is the largest compartment of the eukaryotic cell’s nucleus, known to act as a ribosome factory, thereby sustaining the translation machinery for protein synthesis (Gupta and Santoro, 2020). Increasing evidence indicates that the role of the nucleolus and rRNA genes might go beyond the control of ribosome biogenesis. One such role is linked to the organization of the genome since repressive chromatin domains can often be found in the vicinity of the nucleolus. However, this aspect of genome compartmentalization in the cell’s nucleus has so far remained under-investigated due to technical challenges in the identification of NADs.

In this work, we established novel methods that allowed accurate genome-wide identification of NADs. While Nucleolar-DamID identified all NADs using as readout DNA interactions with the engineered nucleolar histone H2B, HiC-rDNA detected a subclass of NADs that interact with rRNA genes. Although these methods were based on completely different methodologies, they both revealed similar features of chromosome organization around the nucleolus in ESCs and NPCs. Previous methods for NAD identification used sonication-based biochemical purification of nucleoli, a methodology that is subjected to a certain variation in nucleoli preparation between different cell types and relatively biased toward sonication-resistant heterochromatin (Bizhanova et al., 2020; Dillinger et al., 2017; Lu et al., 2020; Nemeth et al., 2010; van Koningsbruggen et al., 2010; Vertii et al., 2019). Relative to this methodology, the application of Nucleolar-DamID and HiC-rDNA has no bias for any kind of chromatin state and does not vary between cell types, thereby allowing direct comparisons of chromosome architecture around the nucleolus between distinct cell states. We found some similarities between NAD maps obtained with previous nucleoli purifications and our methodologies, such as repressive chromatin features and enrichment in olfactory receptor and zinc finger genes (Nemeth et al., 2010; van Koningsbruggen et al., 2010). It is important here to note that major and minor satellites, which composed centric and pericentric domains and known to be in contact with the nucleolus, could not be identified by Nucleolar-DamID due to the lack of GATC motif. However, Nucleolar-DamID and HiC-rDNA revealed further and novel details in chromosome architecture around the nucleolus that did not emerge from previous studies. Further, NAD maps provided in this work can be integrated in future studies of 3D genome organization that have so far excluded the nucleolus because of the lack of profiles of genome interactions with the nucleolus.

The analysis of NADs in ESCs revealed that 50% of NADs coincide with LADs, a result that is consistent with previous data showing that chromosomal regions that were LADs in the mother cell could be repositioned close to the nucleolus after completion of cell division (Kind et al., 2013). This work not only identified these LADs able to contact the nucleolus but also determined that these LADs differ in the chromatin composition relative to LADs that are excluded from the nucleolus (LAD-only). Thus, the identification of NADs revealed further layers of repressive compartments, showing distinct chromatin features between genomic domains positioned closed to the nucleolus, NL, or both. Indeed, although NADs shared some similarity with known features of LADs, such as low overall gene density and low expression levels (van Steensel and Belmont, 2017), they also displayed some distinct characteristic. In particular, NAD/LAD demarcate regions of the genome that are enriched in H3K9me2 and depleted in H3K27me3 relative to LAD-only. This was also evident by the depletion of Polycomb components Ezh2 and Ring1b at the NAD borders whereas LAD-only were enriched as previously reported (Harr et al., 2015). Thus, contacts with the nucleolus, even for sequences able to interact with the NL, shape a distinct repressive chromatin state.

The enrichment of NADs for H3K9me2 was of particular interest since previous work linked this modification with the chromatin state of rRNA genes (Savić et al., 2014). The induction of repressive chromatin at rRNA genes in ESCs was shown to cause global genome remodelling toward a more heterochromatic state, including the formation of highly condensed heterochromatic structures (Savić et al., 2014). In this work, we showed that the repressive chromatin of rRNA genes induced the increase of H3K9me2 at genomic regions adjacent to NADs whereas regions neighboring LAD-only were not affected. These results support a role of the nucleolus in establishing repressive chromatin states at regions close to the nucleolus. It is important here to note that in ESCs all rRNA genes are transcriptionally and epigenetically active compared to differentiated cells and this state is required to maintain pluripotency (reviewed in (Gupta and Santoro, 2020). However, this hyperactive state of rRNA genes in ESCs does not positively correlate with elevated protein synthesis relative to differentiated cells (Guzzi et al., 2018; Ingolia et al., 2011; Sampath et al., 2008). Further, upon differentiation, a fraction of rRNA genes acquire repressive chromatin features, a process that is required to exit from pluripotency (Leone et al., 2017) and timely coincident with the global remodelling of the genome toward a repressed state (Efroni et al., 2008). The results of this work strongly suggest that *de novo* silencing of rRNA gene during ESC differentiation might increase the concentration of repressive chromatin regulators in the nucleolus and serve to establish repressive states at genomic regions neighbouring the nucleolus, thereby linking nucleolar and nuclear chromatin states.

The analysis of NAD composition during differentiation of ESCs into NPCs revealed a common, global chromosome architecture around the nucleolus of both cell types. However, there were some remarkable differences. Compared to ESCs, the structures of all chromosomes around the nucleolus of NPCs appears more compact and rigid, displaying fewer unique contacts but with higher interaction frequency, in particular at centromere-proximal regions. These results are consistent with previous reports showing that ESCs harbor a more open and dynamic chromatin than differentiated cells (Gaspar-Maia et al., 2011; Meshorer et al., 2006). Further, we found cell type specific NADs, unique to ESC or NPCs. Regions detaching from the nucleolus increased the interaction strength within the active A compartment whereas contacts in the repressive B compartment decreased, underscoring the nucleolus as a scaffold for repressive chromatin domains. However, moving away from the nucleolus did not generally correspond to gene activation. Considering that genes detached from the nucleolus of NPCs were highly related to neurogenesis, it appears that the release from the nucleolus might unlock these genes for activation for a later stage of differentiation. A similar observation was also reported for LADs (Peric-Hupkes et al., 2010). Consistent with these results, genes that relocated away from nucleoli of ESCs to NL in NPCs, were related to pathways that should not be activated during neurogenesis, thereby remaining in contact with the other repressive compartment.

In summary, our work established novel methodologies to identify NADs in every cell type and allowed to finally distinguish distinct layers of organization of the genome at repressive compartments that depend on the interaction with the nucleolus, NL, or both. We predict that the application of Nucleolar-DamID and HiC-rDNA will be relevant for future works, including the understanding of the determinants for specific targeting to nucleolus or NL or the application of single cell analyses that were prohibitive with the previous biochemical based-methods. Further, considering that structural changes in the nucleolus are often observed in cancer and premature ageing (Buchwalter and Hetzer, 2017; Hein et al., 2013; Weeks et al., 2019), the identification and analysis of NAD composition will be important for the understanding of alterations of 3D genome organization in disease linked to nucleolus alterations. Finally, the identification of NADs by Nucleolar-DamID will feed the study of genome organization and provide novel insights into basic principles of genome organization and its role in gene expression and cell function.

## Acknowledgments

This work was supported by the Swiss National Science Foundation (31003A_173056 to R.S), and ERC grant (ERC-AdG-787074-NucleolusChromatin). We thank Peter Meister for the assistance in DamID analysis, Catherine Aquino and the Functional Genomic Center Zurich for the assistance in sequencing, and the Center for Microscopy and Image Analysis of the University of Zurich.

## Contributions

C.B. and D.B. cloned the constructs for the nucleolar DamID. CB established ESC lines, set the conditions for the Nucleolar-DamID, performed experiments and data analysis of NADs. J.J.-R. performed and analyzed H3K9me2 ChIP-seq experiments. S.G. performed and analyzed FISH experiments. R.K. analyzed Hi-C data. All authors contributed to experimental design and data interpretation. R.S conceived and supervised the project.

## Accession numbers

All raw data generated in this study using high throughput sequencing are accessible through NCBI’s GEO (accession number GSE150822).

## Material and Methods

### Cell culture

One hundred and twenty-nine mouse embryonic stem cells (E14 line) were cultured in either 2i media composed of DMEM-F12 and Neurobasal medium (1:1, Life Technologies), supplemented with 1× N2/B27 (Life Technologies), 1× penicillin/streptomycin/l-glutamine (Life Technologies), 50 μM β-mercaptoethanol (Life Technologies), recombinant leukemia inhibitory factor, LIF (Polygene, 1,000 U/ml) and MEK and GSK3β inhibitors, 2i (Sigma CHIR99021 and PD0325901, 3 and 1 μM, respectively). ESCs were seeded at a density of 50,000 cells/cm^2^ in culture dishes (Corning^®^ CellBIND^®^ surface) coated with 0.1% gelatin without feeder layer. Propagation of cells was carried out every 2 days using enzymatic cell dissociation.

NIH3T3 and HEK 293T cells were cultured in Dulbecco’s modified Eagle’s medium (DMEM, Life Technologies) supplemented with 10% fetal calf serum (FCS, Biowaste) and 1% penicillin/streptomycin (Life Technologies).

Neural progenitor cells were generated from ESCs, according to a previously established protocol (Bibel et al., 2004) In brief, differentiation used a suspension-based embryoid bodies formation (Bacteriological Petri Dishes, Bio-one with vents, Greiner). The neural differentiation media (DMEM, 10% fetal calf serum, 1× MEM NEAA, 2 mM Pen/Strep, β-mercaptoethanol, and sodium pyruvate) was filtered through 0.22 μm filters and stored at 4°C. During the 8-day differentiation procedure, media was exchanged every 2 days. In the last 4 days of differentiation, the media was supplemented with 2 μM retinoic acid to generate neural precursors that are Pax-6-positive radial glial cells.

### Establishment of ESC lines for Nucleolar-DamID

CRISPR/Cas9 cloning and targeting strategy was performed as previously described (Ran et al., 2013). ESCs were co-transfected with a plasmid expressing the Cas9 proteins and the sgRNA guide sequence targeting the *Rosa26* locus (Genome CRISPR™ mouse ROSA26 safe harbor gene knock-in kit, SH054, GeneCopoeia) and the HDR repair template plasmid containing either H2B-Dam or H2B-Dam-NoLS constructs flanked by the homology arms with a molar ratio of 1:3.

Two days after transfection, ESCs were selected using 2ug of Puromycin (Life Technologies) overnight. After recover, ESCs were further treated with 1ug of Puromycin (Life Technologies) for three days. After additional three days of recover, cells were seeded for single cell clone isolation. Resistant ESC clones were genotyped by PCR using primers able to distinguish between insertions of the construct in one or both alleles.

### Transfections

1.5 × 10^5^ NIH3T3 cells were plated in 6-well plates and transfected respectively with 200ng of plasmid expressing H2B-GFP or H2B-GFP-NoLS under the control of minimal CMV promoter, or with 1μg of plasmid expressing TTFI-GFP under the full CMV promoter using Transit-X2 transfection reagent (Mirus) in Opti-MEM GlutaMAX reduced-serum medium (Life Technologies). When indicated, 24h post-transfection, cells were treated with 50 ng of Actinomycin D (Sigma) and the other half were refreshed with new media. 24h later, cells were fixed and mounted with DAPI-mounting media (Vector) for imaging.

2 × 10^5^ HEK 293T cells were plated in 6-well plates. The day after, cells were transfected with 500 ng of plasmid containing the sequence *hsp-TetO-GFP-Dam-DD-EF1a-puro-T2A-TetR-KRAB* using calcium phosphate protocol. 24h post-transfection, part of the cells were induced with 1μg/mL doxycycline and 1μM Shield1, and the remaining cells were refreshed with new media. The day after the treatment, cells were imaged under the microscope.

9 × 10^5^ ESC were plated in gelatin-coated 10cm culture dish and transfected with 42 μg pRNA, or RNA-control using Lipofectamine MessengerMAX reagent (Invitrogen) in Opti-MEM GlutaMAX (Life Technologies) reduced-serum medium. 48h post-transfection, ESCs were collected for downstream analyses.

### Immunofluorescence

Live cell images of NIH3T3 cells (**Fig. 1B**) and HEK 293T cells **(Fig. S1B**) were taken and digitally recorded using FLoid Cell Imaging Station (ThermoFisher). In the experiments with ActD treatments (**Fig. 1D**), cells were grown on glass coverslips, fixed with 4% paraformaldehyde, and stained with DAPI. Images were taken and digitally recorded using Leica DMI6000 B microscope.

### Chromatin fractionation

48 hours post-transfection, NIH3T3 cells were collected by trypsinization, washed once with PBS and counted. ES cell pellets were resuspended at a concentration of 10mio cells/ml in chromatin fractionation buffer (10mM Hepes pH 7.6, 150mM NaCl, 3mM MgCl_2_, 0.5% Triton X-100, 1mM DTT freshly supplemented with cOmplete^™^ Protease Inhibitor Cocktail (Roche)) and incubated for 30 minutes at room temperature rotating. Precipitated chromatin was fractionated by centrifugation. Total and chromatin fractionated samples were further processed by MNase (S7 Micrococcal nuclease, Roche) digest for ensuring sufficient genomic DNA fragmentation. All samples were incubated in 1x Laemmli buffer (10% glycerol, 10mM Tris pH 6.8, 2% SDS, 0.1mg/ml bromophenolblue, 2% β-mercaptoethanol) at 95°C for 5 minutes and were further analyzed by Western Blotting.

### DNA-FISH

DNA fluorescence in situ hybridization (DNA-FISH) was performed with Agilent SureFISH DNA-FISH probes following the manufacturer’s protocol with modifications as mentioned below. ESCs were cultured on matrigel coated coverslips. The coverslips were fixed using 3.7% Formaldehyde for 10 min at room temperature. Cells were washed twice with 1X PBS and then dehydrated with graded ethanol concentrations up to 100% ethanol and air dried. The coverslips were incubated with 10 μL mixture of a custom probe set targeting a selected DNA locus (Agilent) and SureFISH Hybridization Buffer (Agilent, G9400A) with turned cell-side down. The coverslips and probe mixture were denatured for 8 min at 83 °C, then incubated at 37 °C overnight in a dark humidified chamber. Next day, coverslips were washed with FISH Wash Buffer 1 (Agilent, G9401A) at 73 °C for 2 min on a shaking incubator at 300 rpm, and FISH Wash Buffer 2 (Agilent, G9402A) at room temperature for 1 min. Following DNA-FISH probe hybridization, immunofluorescence was performed. Coverslips were prepared for immunofluorescence by rehydration and suspenion in 1X PBS. Coverslips were permeabilized with 0.5% Triton-X in 1X PBS on ice for 5 minutes followed by incubation at room temperature for 10 min, and kept in blocking buffer (1%BSA in 1XPBS-Tween 20 (0.1%) at room temperature for 1 hr. Coverslips were then incubated with primary anti-Nucleolin antibodies (Abcam; ab22758; 1:100) in a humidified chamber overnight at 4°C. After washing with 0.1% Triton-X in 1XPBS at room temperature, they were incubated with secondary antibodies in a dark humidified chamber at room temperature for 1.5 hour. After washing with 1X PBS buffer, they were stained with Hoechst 33342 followed by mounting. The secondary antibodies used for IF were goat anti-rabbit IgG (H+L) highly cross-adsorbed secondary antibody, Alexa Fluor 488 (Thermo Fisher Scientific; A11034; 1:500) and goat anti-rabbit IgG (H+L) highly cross-adsorbed secondary antibody, Alexa Fluor 546 (Thermo Fischer Scientific; A11035; 1:500). Probe sets used for FISH were designed by Agilent technologies using their standard procedures against genomic regions defined in **Table S12**. DNA FISH/IF samples were imaged using a Leica SP8 upright Microscope, with a z-stack collected for each channel (step size, 0.15 um, frame interval 1 sec). The objective used was HC PL APO CS2 63x/1.40 oil objective. Image were processed by ImageJ 1.52p. The individual cells were identified by Hoechst staining and cells containing signal for DNA-FISH channel were identified manually on the corresponding fluorescent channel (with background subtraction). Images were cropped to contain single cell.

### RNA extraction, reverse transcription, and quantitative PCR (RT-qPCR)

RNA was purified with TRIzol reagent (Life Technologies). 1μg total RNA was primed with random hexamers and reverse-transcribed into cDNA using MultiScribe™ Reverse Transcriptase (Life Technologies). Amplification of samples without reverse transcriptase assured absence of genomic or plasmid DNA (data not shown). The relative transcription levels were determined by normalization to *beta-Actin* mRNA levels, as indicated. qRT-PCR was performed with KAPA SYBR^®^ FAST (Sigma) on a Rotor-Gene Q (Qiagen).

### In vitro synthesis of pRNA

Topo2.1 plasmids with insertion of pRNA and RNA-control sequences were previously described (Savić et al., 2014). BamHI-linearized plasmids were *in vitro* transcribed with T7 RNA polymerase (Thermo Fisher EP0111). Synthesized RNA transcripts were verified by agarose gel electrophoresis and purified using NucleoSpin RNA II column (Machere-Nagel, cat. no. 740955) according to manufacturer’s protocol.

### DamIDseq

7 × 10^4^ H2B-Dam and H2B-Dam-NoLS ESC lines were seeded in 6-well plates (Corning^®^ CellBIND^®^ surface) coated with 0.1% gelatin without feeder layer. to induce the expression of Dam-fused proteins, two days after seeding, half of the cells were treated with 100ng/ml Doxycycline and 1μM Shield1 (Clontech Takara) for 15h. For the analysis of NPCs, 8 days after differentiation, H2B-Dam and H2B-Dam-NoLS cells were harvested with 10X trypsin to get single cells from the embryoid bodies. 1 × 10^6^ cells NPCs were seeded in 6-well plates coated with 0.1% gelatin and treated with 100ng/ml Doxycycline and 1μM Shield1 for 15h. The day after, cells were harvested by trypsinization and DNA was extracted using Quick-DNA Miniprep Plus kit (Zymo Research). To test the efficiency and the specificity of the treatment, quantitative measurements of the m6A levels at GATC of rRNA genes and *Tuba1a* were assessed by Dpn II digestion followed by qPCR as previously described (Kind et al., 2015).

DamID-seq was performed using previously described protocols (Vogel et al., 2007). Briefly, 500ng of genomic DNA was digested for 4h at 37°C with Dpn I (New England Biolabs) to cut methylated GATC sites. After heat inactivation for 20 minutes at 80°C, DamID adaptors were blunt-ended and ligated overnight at 16°C, followed by heat inactivated at 65°C for 10 minutes. In order to cut unmethylated GATC sequences, DNA was digested for 3h at 37°C with Dpn II (New England Biolabs) and heat inactivated at 65°C for 20 minutes. Adaptor-ligated fragments were amplified using the Advantage® GC 2Polymerase mix (Clontech Takara) (primer described in Table MM, Vogel et al., 2007). DamID libraries were purified using Agencourt AMPure XP beads (Beckam Coulter). Since the size of fragments produced exceeded the one fitting the sequencing machine, the libraries were fragmented using the ds Fragmentase (New England Biolabs) to enrich the concentration of the libraries below 500bp and, afterwards, the libraries were purified again using the Agencourt AMPure XP beads (Beckam Coulter). The quantity and quality of the isolated DNA was determined with a Qubit® (1.0) Fluorometer (Life Technologies, California, USA). The Nugen Ovation Ultra Low Library Systems (Nugen, Inc, California, USA) was used to prepare the libraries for Illumina sequencing. Briefly, Nucleolar-DamID samples (1 ng) were end-repaired and polyadenylated. Then, Illumina compatible adapters, containing the index for multiplexing, were ligated. The quality and quantity of the enriched libraries were validated using Qubit® (1.0) Fluorometer and the Bioanalyzer 2100 (Agilent, Waldbronn, Germany). The libraries were buffered tin 10nM in Tris-Cl 10 mM, pH8.5 with 0.1% Tween 20. The TruSeq SR Cluster Kit v4-cBot-HS (Illumina, Inc, California, USA) was used for cluster generation using 8 pM of pooled normalized libraries on the cBOT. Using the TruSeq SBS Kit v4-HS (Illumina, Inc, California, USA) the sequencing was performed as pair-end 150bp reads using the Illumina NovaSeq 6000. Sequences are aligned to the mouse reference genome mm10 using Bowtie2 (version 2.3.4.3) (Langmead and Salzberg, 2012). Resulting sam files were converted into bam files, sorted and indexed using samtools (version 1.9)(Li et al., 2009). The bam files were analyzed using the damidseq pipeline script from the Brand group (http://owenjm.github.io/damidseq_pipeline) using a 100kbp resolution(Marshall and Brand, 2015). The pipeline bins the mapped reads into GATC-fragments according to GATC-sites indicated by a gff file for the GRCm38 mouse genome (already provided by the authors of the pipeline on the website above) and normalizes reads against the Dam control, in our case the H2B-Dam sample. The pipeline gives a bedgraph file with the log_2_ ratio of the m6A between H2B-Dam-NoLS and the H2B-Dam only. The bedgraphs were visualized using the tool Integrative Genome Viewer (IGV, version 2.5.2)(Robinson et al., 2011) to extract representative Nucleolar-DamID tracks. The beadgraph files were converted into bigwig files using the bedGraphToBigWig UCSC package (version 4) and the Pearson correlation was assessed using “multiBigwigSummary” and “plotPCA” from deepTools (version 3.2.1) (Quinlan and Hall, 2010). The bedgraph files were processed with the find_peaks software associated with the pipeline (https://github.com/owenjm/find_peaks) adjusting the values of the FDR and the minimum quantile (FDR <0.01 and min_quant 0.70). Only the significative peaks common to both replicates were considered as NADs for further analysis.

The identification of NADs overlapping with LADs, genomic contacts with rRNA genes identified by HiC-rDNA, A and B compartment, early/late replicating regions, and the identification of ESC-sp and NPCsp-NADs were performed using “Intersect intervals” from bedtools (version 2.28.0) (Quinlan and Hall, 2010). NAD-only and NAD/LAD distribution over the chromosomes, was generated with ChIPseeker (Yu et al., 2015) in R studio (version 1.0.44). GO term analysis was performed using DAVID 6.8 (Huang et al., 2009).

### ChIPseq

ChIP analysis was performed as previously described (Leone et al., 2017). Briefly, 1% formaldehyde was added to cultured cells to cross-link proteins to DNA. Isolated nuclei were then lysed with lysis buffer (50mM Tris-HCl, pH 8.1, 10 mM EDTA, pH 8, 1% SDS, 1X protease inhibitor cOmplete EDTA-free cocktail, Roche). Nuclei were sonicated using a Bioruptor ultrasonic cell disruptor (Diagenode) to shear genomic DNA to an average fragment size of 200 bp. 20 μg of chromatin was diluted to a total volume of 500 μl with ChIP buffer (16.7 mM Tris–HCl, pH 8.1, 167 mM NaCl, 1.2 mM EDTA, 0.01% SDS, 1.1% Triton X-100) and incubated overnight with the ChIP-grade antibodies against H3K9me2 and H3K9me3. After washing, bound chromatin was eluted with the elution buffer (1% SDS, 100 mM NaHCO_3_). Upon proteinase K digestion (50°C for 3 h) and reversion of cross-linking (65°C, overnight), DNA was purified with phenol/chloroform, ethanol precipitated and quantified by qPCR..

For ChIPseq analyses, the quantity and quality of the isolated DNA was determined with a Qubit® (1.0) Fluorometer (Life Technologies, California, USA) and a Bioanalyzer 2100 (Agilent, Waldbronn, Germany). The Nugen Ovation Ultra Low Library Systems (Nugen, Inc, California, USA) was used in the following steps. Briefly, ChIP samples (1 ng) was end-repaired and polyadenylated before the ligation of Illumina compatible adapters. The adapters contain the index for multiplexing. The quality and quantity of the enriched libraries were validated using Qubit® (1.0) Fluorometer and the Bioanalyzer 2100 (Agilent, Waldbronn, Germany). The libraries were normalized to 10nM in Tris-Cl 10 mM, pH8.5 with 0.1% Tween 20. The TruSeq SR Cluster Kit v4-cBot-HS (Illumina, Inc, California, USA) was used for cluster generation using 8 pM of pooled normalized libraries on the cBOT. Sequencing was performed on the Illumina HiSeq 2500 single end 126 bp using the TruSeq SBS Kit v4-HS (Illumina, Inc, California, USA).

### ChIPseq data analysis

Own and published ChIPseq reads were aligned to the mouse mm10 reference genome using Bowtie2 (version 2.3.4.3)(Langmead and Salzberg, 2012). Read counts were computed and normalized using “bamCoverage” from deepTools (version 3.2.1)(Ramirez et al., 2014) using a bin size of 50bp. To calculate read coverage for 20kb bin region of H3K9me2 and H3K9me3 ChIPseq, “multiBamSummary” from deepTools was used. The border profiles and the read coverage box plots were generated using deepTools. H3K9me2 increase in ESC+pRNA regions distribution over the chromosomes was generated with the ChIPseeker package(Yu et al., 2015). Integrative Genome Viewer (IGV, version 2.5.2)(Robinson et al., 2011) was used to visualize and extract representative ChIPseq tracks.

### Identification of rDNA contacts by Hi-C

The identification of genomic contacts with rRNA genes was performed by recovering reads containing rRNA gene contacts from three published Hi-C data of ESCs and NPCs (Bonev et al., 2017). The obtained Hi-C data sets have been analyzed with Juicer (Durand et al., 2016) all in one computational pipeline for generating Hi-C maps from raw fastq data files and command line tools for feature annotation on the Hi-C maps. During Juicer analysis, raw fastq data sets have been aligned to the customized mm10 genome with Burrows-Wheeler Aligner (Li and Durbin, 2009) under default parameters. The modified mm10 genome contained one rRNA gene unit attached to the end of chromosome 12. The chromosomal interaction have been extracted from interaction matrices (hic files) with Juicebox tools command *dump* under the following parameters (contacts: observed, normalization applied: Knight-Ruiz matrix balancing(Knight and Ruiz, 2012) under base-pair delimited resolution with bin size 5000). ENCODE Data Analysis Consortium Blacklisted Regions(Hoffman et al., 2013) were excluded from the analysis with bedtools (Quinlan and Hall, 2010). Only Hi-C reads contacting rRNA gene sequences and other genomic sequences have been selected for further analysis through the computation with bedtools pairtoBED function(Quinlan and Hall, 2010) and Python Pandas Library (https://pandas.pydata.org/). Common contacts between the three Hi-C replicates were identified using HiCcompare (Stansfield et al., 2018), running the tool under default parameters.

### RNAseq

Total RNA was purified with TRIzol reagent (Life Technologies). The quality of the isolated RNA was determined with a Qubit® (1.0) Fluorometer (Life Technologies, California, USA) and a Fragment Analyzer (Agilent, Santa Clara, California, USA). Only those samples with a 260 nm/280 nm ratio between 1.8–2.1 and a 28S/18S ratio within 1.5-2 were further processed. The TruSeq Stranded mRNA (Illumina, Inc, California, USA) was used in the succeeding steps. Briefly, total RNA samples (100-1000 ng) were polyA enriched and then reverse-transcribed into double-stranded cDNA. The cDNA samples was fragmented, end-repaired and adenylated before ligation of TruSeq adapters containing unique dual indices (UDI) for multiplexing. Fragments containing TruSeq adapters on both ends were selectively enriched with PCR. The quality and quantity of the enriched libraries were validated using Qubit® (1.0). The product is a smear with an average fragment size of approximately 260 bp. Libraries were normalized to 10nM in Tris-Cl 10 mM, pH8.5 with 0.1% Tween 20. The HiSeq 4000 (Illumina, Inc, California, USA) was used for cluster generation and sequencing according to standard protocol. Sequencing were paired end at 2 X150 bp or single end 100 bp. The quality of the 120 bp single end reads generated by the machine was checked by FastQC, a quality control tool for high throughput sequence data (Andrews, 2010). The quality of the reads was increased by applying: a) SortMeRNA (Kopylova, 2012) (version 2.1) tool to filter ribosomal RNA; b) Trimmomatic (Bolger, 2014) (version 0.36) software package to trim the sorted (a) reads. The sorted (a), trimmed (b) reads were mapped against the mouse genome (mm10) using the default parameters of the STAR (Spliced Transcripts Alignment to a Reference, version 2.4.0.1) (Dobin et al., 2013). For each gene, exon coverage was calculated using a custom pipeline and then normalized in reads per kilobase per million (RPKM) (Mortazavi et al., 2008), the method of quantifying gene expression from RNA sequencing data by normalizing for total read length and the number of sequencing reads.

## Supplementary Figures

**Figure S1.**
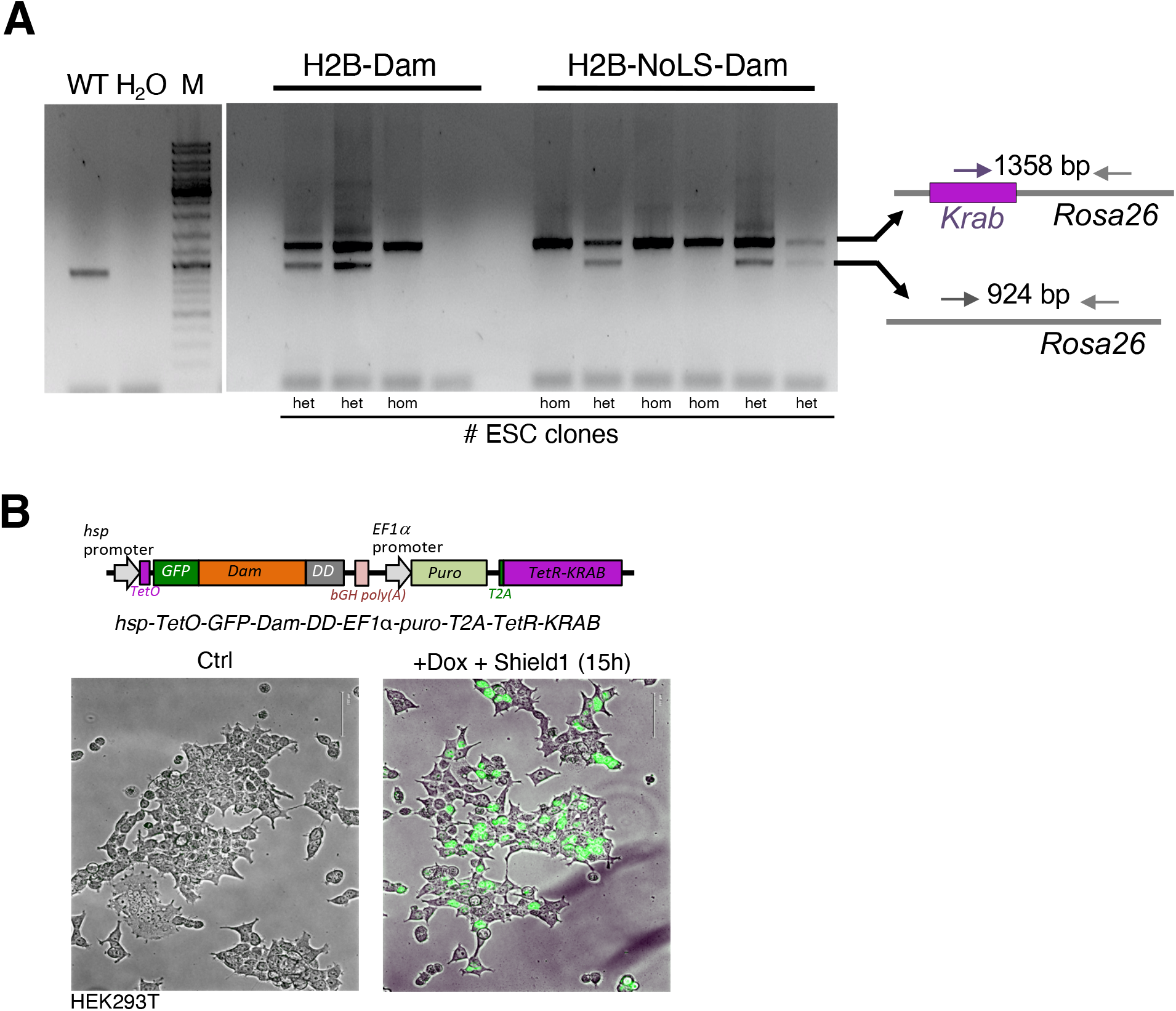
Establishment of Nucleolar-DamID. **A.** PCR genotyping of ESC clones for insertion of the H2B or nucleolar H2B sequences into *Rosa26* locus. het.: heterozygotic insertion; hom.: homozygotic insertion; WT: wild type ESCs; M: DNA marker. **B.** Life cell images showing GFP signal in HEK293T transfected with the *hsp-TetO-GFP-Dam-DD-EF1*α*-puro-T2A-TetR-KRAB* plasmid and treated without or with 1 μg/ml doxycycline (Dox) and 1 μM Shield1 for 15 hours.

**Figure S2.**
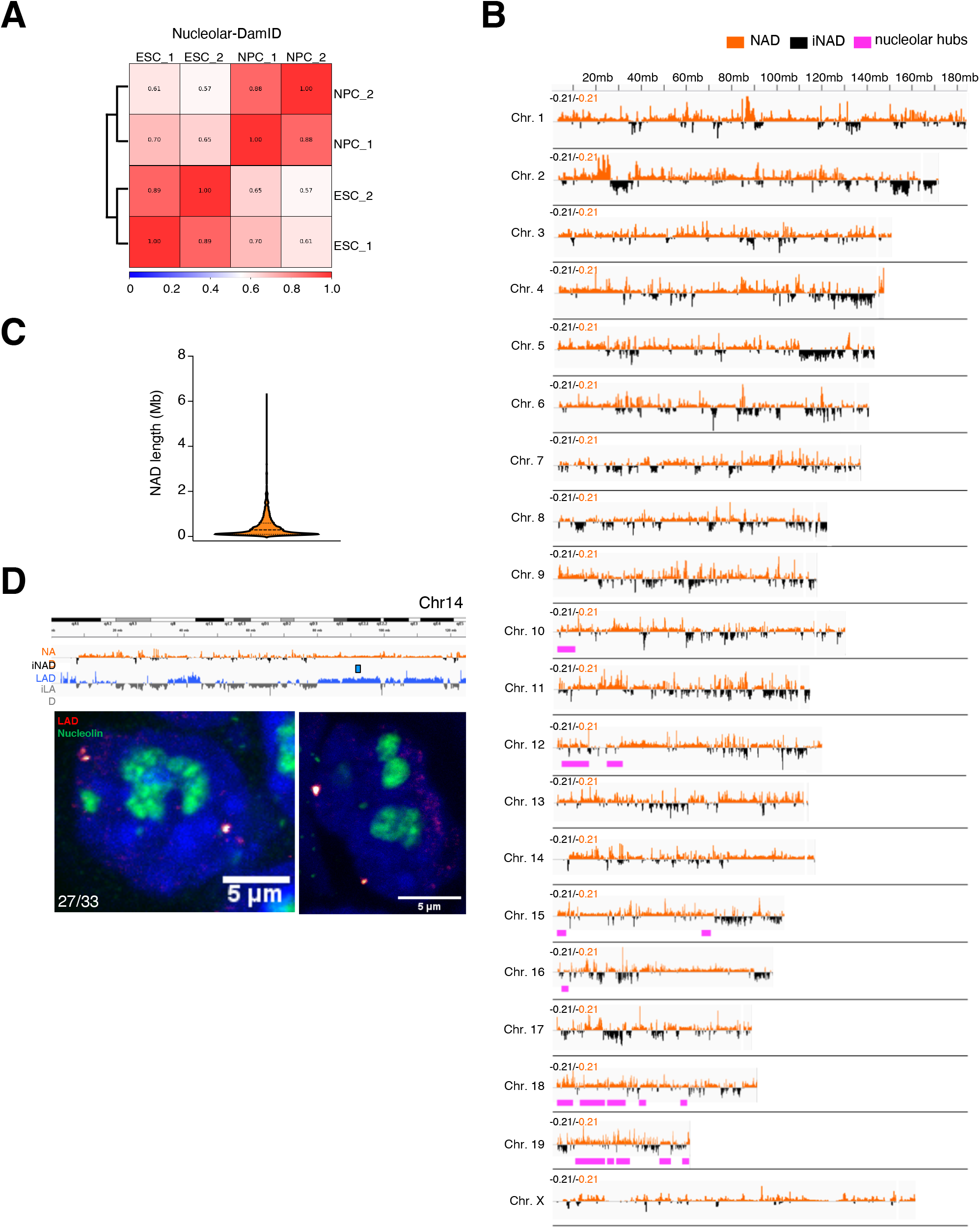
Nucleolar-DamID in ESCs. **A.** Pearson correlation of Nucleolar-DamID experiments in ESCs and NPCs. **B.** Violin plot showing distribution of NAD length. **C.** Chromosomal view of NADs in ESCs. NADs are measured as log_2_ ratio of m6A levels between H2B-Dam-NoLS vs. H2B-Dam. iNAD: inter-NAD regions. iLAD: inter-LAD regions. Pink bars correspond to sequence found in nucleolar hubs using SPRITE method (Quinodoz et al., 2018). **D.** Upper panel. NAD and LAD composition of chromosome 14 and FISH probe corresponding to a strong LAD (blue bar). Lower panel. Representative images from immunofluorescence for nucleolin (green) combined with DNA FISH using probes corresponding to LAD-only region of chromosome 14 (red), and DAPI (blue). The signal of the LAD probe was located close to NL in the majority of analyzed ESCs (27/33).

**Figure S3.**
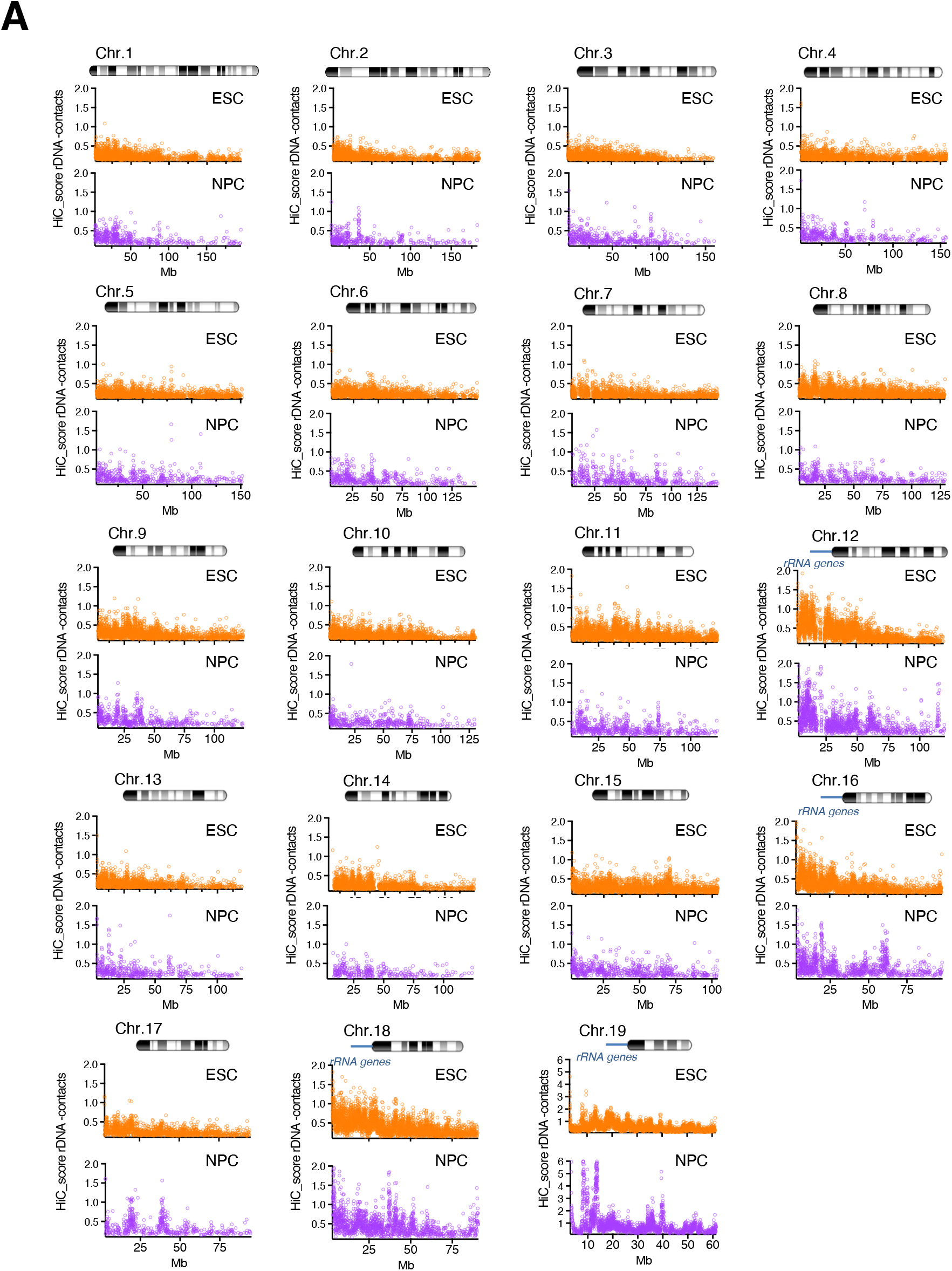
rDNA contacts identified by HiC-rDNA in ESCs and NPCs. **A.** Count score of rDNA contacts obtained by HiC-rDNA on chromosomes of ESCs (orange) and NPCs (magenta). Chromosomes containing rRNA genes (12, 16, 18, and 19) are indicated.

**Figure S4.**
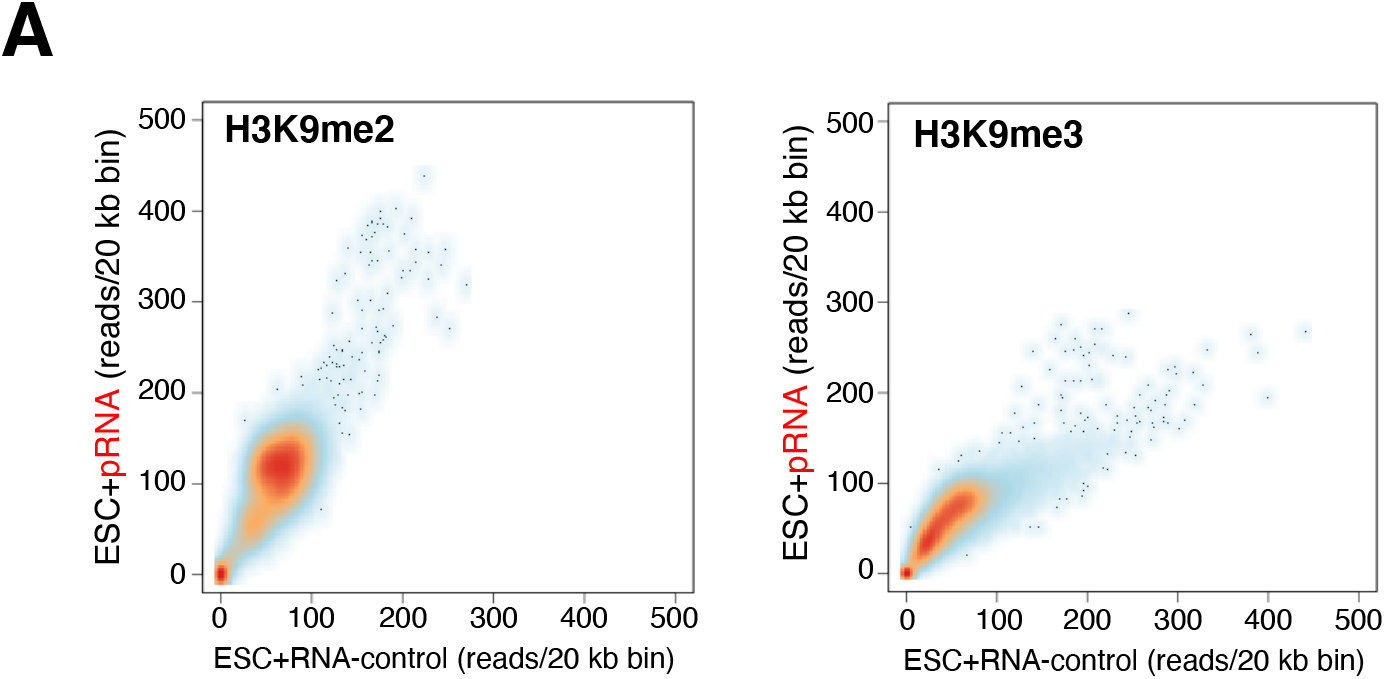
pRNA-mediated heterochromatin formation at rRNA genes increases H3K9me2 at sequences adjacent to the nucleolus. **A.** Addition of pRNA in ESCs caused an increase in H3K9me2 at several genomic regions. Independent ChIPseq experiment of ESCs transfected with pRNA and RNA control. Scatter plot showing H3K9me2 and H3K9me3 levels (reads/20kb bin) between ESC+pRNA and ESC+RNA-control.

**Figure S5.**
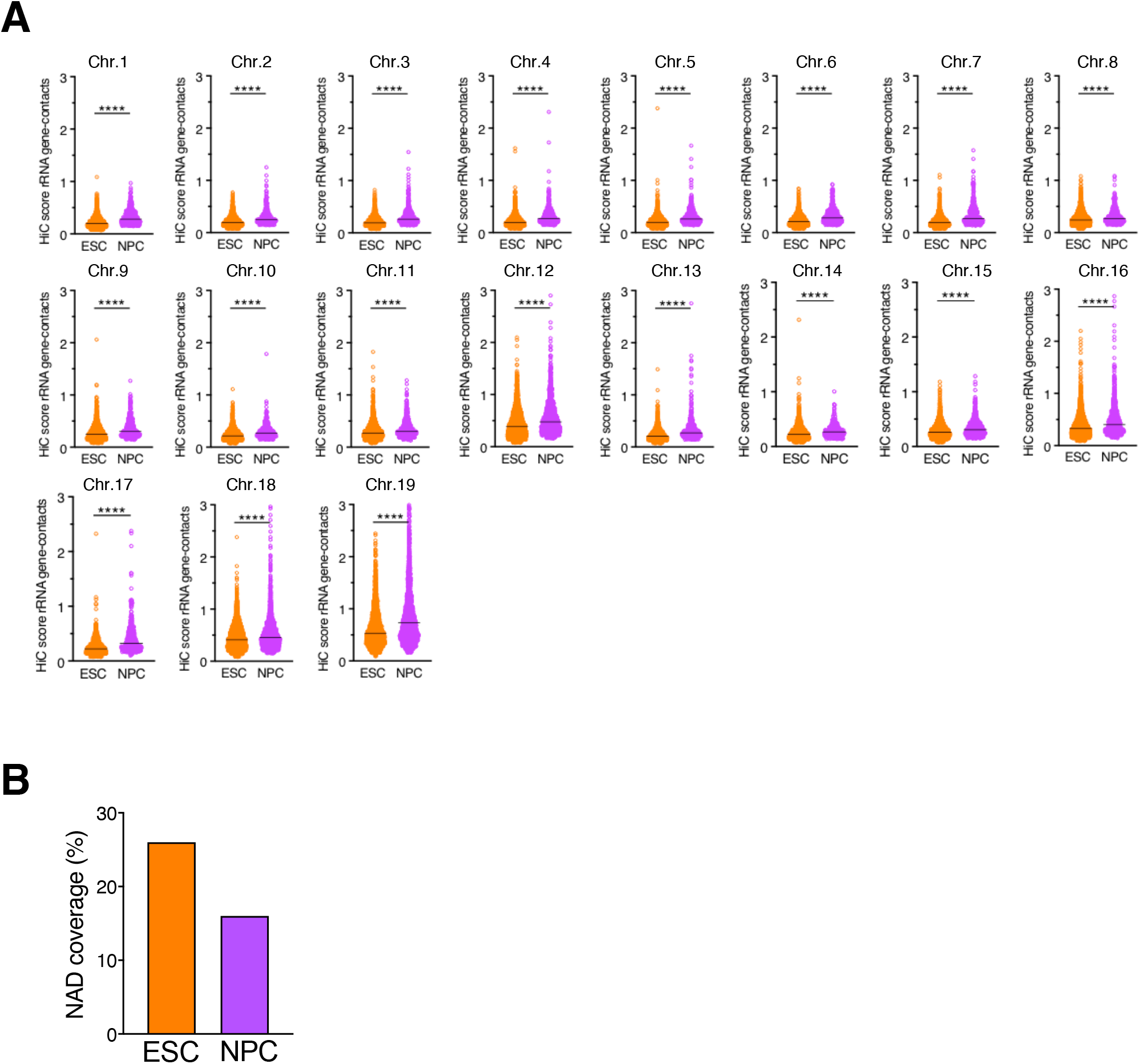
rDNA contacts in ESCs and NPCs. **A.** Count score of rDNA contacts at each chromosome in ESCs (orange) and NPCs (magenta). Statistical significance (*P*-values) between ESCs and NPCs was calculated using the paired two-tailed t-test ****< 0.0001).

**Figure S6.**
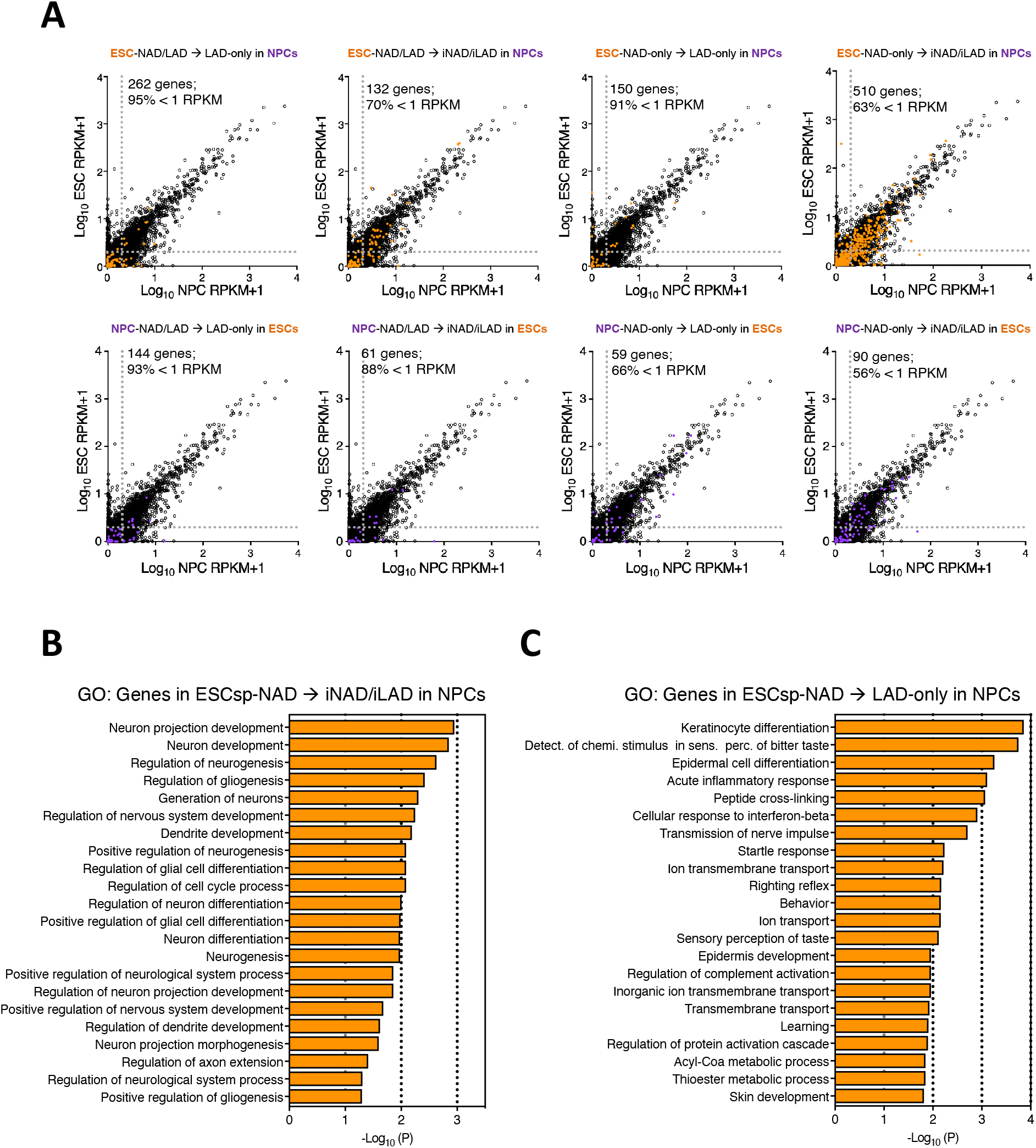
ESC and NPC specific NADs. **A.** Scatter plot showing gene expression levels between ESC and NPCs. Expression of genes located at ESC_sp_- and NPC_sp_-NAD types (NAD/LAD and NAD-only) and their location in NPCs and ESCs, respectively, is shown. Genes located at ESC_sp_- and NPC_sp_-NAD are highlighted in orange and magenta, respectively, whereas total genes are represented in black. Dotted lines indicate RPKM value as 1. **B,C.** Gene ontology terms of genes located at ESC_sp_-NAD that are located at iNAD/iLAD (**A**) and LAD-only (**B**) in NPCs.

## References

Andersen, J.S., Lam, Y.W., Leung, A.K., Ong, S.E., Lyon, C.E., Lamond, A.I., and Mann, M. (2005). Nucleolar proteome dynamics. Nature 433, 77–83.

Andrews, S. (2010). FastQC: a quality control tool for high throughput sequence data.

Aughey, G.N., Estacio Gomez, A., Thomson, J., Yin, H., and Southall, T.D. (2018). CATaDa reveals global remodelling of chromatin accessibility during stem cell differentiation in vivo. Elife 7.

Banaszynski, L.A., Chen, L.C., Maynard-Smith, L.A., Ooi, A.G., and Wandless, T.J. (2006). A rapid, reversible, and tunable method to regulate protein function in living cells using synthetic small molecules. Cell 126, 995–1004.

Becker, J.S., McCarthy, R.L., Sidoli, S., Donahue, G., Kaeding, K.E., He, Z., Lin, S., Garcia, B.A., and Zaret, K.S. (2017). Genomic and Proteomic Resolution of Heterochromatin and Its Restriction of Alternate Fate Genes. Mol Cell 68, 1023–1037 e1015.

Belmont, A.S. (2002). Mitotic chromosome scaffold structure: new approaches to an old controversy. Proc Natl Acad Sci U S A 99, 15855–15857.

Bersaglieri, C., and Santoro, R. (2019). Genome Organization in and around the Nucleolus. Cells 8(6), 579.

Bibel, M., Richter, J., Schrenk, K., Tucker, K.L., Staiger, V., Korte, M., Goetz, M., and Barde, Y.A. (2004). Differentiation of mouse embryonic stem cells into a defined neuronal lineage. Nat Neurosci 7, 1003–1009.

Birbach, A., Bailey, S.T., Ghosh, S., and Schmid, J.A. (2004). Cytosolic, nuclear and nucleolar localization signals determine subcellular distribution and activity of the NF-kappaB inducing kinase NIK. J Cell Sci 117, 3615–3624.

Bizhanova, A., Yan, A., Yu, J., Zhu, L.J., and Kaufman, P.D. (2020). Distinct features of nucleolus-associated domains in mouse embryonic stem cells. Chromosoma.

Bolger, A., M.; Lohse, M.; Usadel, B. (2014). Trimmomatic: A flexible trimmer for Illumina Sequence Data. Bioinformatics.

Bonev, B., Mendelson Cohen, N., Szabo, Q., Fritsch, L., Papadopoulos, G.L., Lubling, Y., Xu, X., Lv, X., Hugnot, J.P., Tanay, A., et al. (2017). Multiscale 3D Genome Rewiring during Mouse Neural Development. Cell 171, 557–572 e524.

Buchwalter, A., and Hetzer, M.W. (2017). Nucleolar expansion and elevated protein translation in premature aging. Nat Commun 8, 328.

Dalcher, D., Tan, J.Y., Bersaglieri, C., Peña-Hernández, R., Vollenweider, E., Zeyen, S., Schmid, M.W., Bianchi, V., Butz, S., Roganowicz, M., et al. (2020). BAZ2A safeguards genome architecture of ground-state pluripotent stem cells. EMBO J, in press.

Dekker, J., and Misteli, T. (2015). Long-Range Chromatin Interactions. Cold Spring Harb Perspect Biol 7, a019356.

Desjardins, R., Smetana, K., Steele, W.J., and Busch, H. (1963). Isolation of Nucleoli of the Walker Carcinosarcoma and Liver of the Rat Following Nuclear Disruption in a French Pressure Cell. Cancer Res 23, 1819–1823.

Dillinger, S., Straub, T., and Nemeth, A. (2017). Nucleolus association of chromosomal domains is largely maintained in cellular senescence despite massive nuclear reorganisation. PLoS One 12, e0178821.

Dobin, A., Davis, C.A., Schlesinger, F., Drenkow, J., Zaleski, C., Jha, S., Batut, P., Chaisson, M., and Gingeras, T.R. (2013). STAR: ultrafast universal RNA-seq aligner. Bioinformatics 29, 15–21.

Durand, N.C., Shamim, M.S., Machol, I., Rao, S.S., Huntley, M.H., Lander, E.S., and Aiden, E.L. (2016). Juicer Provides a One-Click System for Analyzing Loop-Resolution Hi-C Experiments. Cell Syst 3, 95–98.

Efroni, S., Duttagupta, R., Cheng, J., Dehghani, H., Hoeppner, D.J., Dash, C., Bazett-Jones, D.P., Le Grice, S., McKay, R.D., Buetow, K.H., et al. (2008). Global transcription in pluripotent embryonic stem cells. Cell Stem Cell 2, 437–447.

Emmott, E., and Hiscox, J.A. (2009). Nucleolar targeting: the hub of the matter. EMBO Rep 10, 231–238.

Evers, R., and Grummt, I. (1995). Molecular coevolution of mammalian ribosomal gene terminator sequences and the transcription termination factor TTF-I. Proc Natl Acad Sci U S A 92, 5827–5831.

Feric, M., Vaidya, N., Harmon, T.S., Mitrea, D.M., Zhu, L., Richardson, T.M., Kriwacki, R.W., Pappu, R.V., and Brangwynne, C.P. (2016). Coexisting Liquid Phases Underlie Nucleolar Subcompartments. Cell 165, 1686–1697.

Gaspar-Maia, A., Alajem, A., Meshorer, E., and Ramalho-Santos, M. (2011). Open chromatin in pluripotency and reprogramming. Nature reviews Molecular cell biology 12, 36–47.

Gonzalez-Sandoval, A., and Gasser, S.M. (2016). On TADs and LADs: Spatial Control Over Gene Expression. Trends Genet 32, 485–495.

Guelen, L., Pagie, L., Brasset, E., Meuleman, W., Faza, M.B., Talhout, W., Eussen, B.H., de Klein, A., Wessels, L., de Laat, W., et al. (2008). Domain organization of human chromosomes revealed by mapping of nuclear lamina interactions. Nature 453, 948–951.

Guetg, C., Scheifele, F., Rosenthal, F., Hottiger, M.O., and Santoro, R. (2012). Inheritance of Silent rDNA Chromatin Is Mediated by PARP1 via Noncoding RNA. Mol Cell 45, 790–800.

Gupta, S., and Santoro, R. (2020). Regulation and Roles of the Nucleolus in Embryonic Stem Cells: From Ribosome Biogenesis to Genome Organization. Stem Cell Reports.

Guzzi, N., Ciesla, M., Ngoc, P.C.T., Lang, S., Arora, S., Dimitriou, M., Pimkova, K., Sommarin, M.N.E., Munita, R., Lubas, M., et al. (2018). Pseudouridylation of tRNA-Derived Fragments Steers Translational Control in Stem Cells. Cell 173, 1204–1216 e1226.

Harr, J.C., Luperchio, T.R., Wong, X., Cohen, E., Wheelan, S.J., and Reddy, K.L. (2015). Directed targeting of chromatin to the nuclear lamina is mediated by chromatin state and A-type lamins. J Cell Biol 208, 33–52.

Hein, N., Hannan, K.M., George, A.J., Sanij, E., and Hannan, R.D. (2013). The nucleolus: an emerging target for cancer therapy. Trends Mol Med 19, 643–654.

Hoffman, M.M., Ernst, J., Wilder, S.P., Kundaje, A., Harris, R.S., Libbrecht, M., Giardine, B., Ellenbogen, P.M., Bilmes, J.A., Birney, E., et al. (2013). Integrative annotation of chromatin elements from ENCODE data. Nucleic Acids Res 41, 827–841.

Huang, W., Sherman, B.T., and Lempicki, R.A. (2009). Systematic and integrative analysis of large gene lists using DAVID bioinformatics resources. Nat Protoc 4, 44–57.

Ingolia, N.T., Lareau, L.F., and Weissman, J.S. (2011). Ribosome profiling of mouse embryonic stem cells reveals the complexity and dynamics of mammalian proteomes. Cell 147, 789–802.

Kempfer, R., and Pombo, A. (2019). Methods for mapping 3D chromosome architecture. Nat Rev Genet.

Kind, J., Pagie, L., de Vries, S.S., Nahidiazar, L., Dey, S.S., Bienko, M., Zhan, Y., Lajoie, B., de Graaf, C.A., Amendola, M., et al. (2015). Genome-wide maps of nuclear lamina interactions in single human cells. Cell 163, 134–147.

Kind, J., Pagie, L., Ortabozkoyun, H., Boyle, S., de Vries, S.S., Janssen, H., Amendola, M., Nolen, L.D., Bickmore, W.A., and van Steensel, B. (2013). Single-cell dynamics of genome-nuclear lamina interactions. Cell 153, 178–192.

Knight, P.A., and Ruiz, D. (2012). A fast algorithm for matrix balancing. IMA Journal of Numerical Analysis 33, 1029–1047.

Kopylova, E.N., L.; Touzet, H. (2012). SortMeRNA: Fast and accurate filtering of ribosomal RNAs in metatranscriptomic data. Bioinformatics.

Langmead, B., and Salzberg, S.L. (2012). Fast gapped-read alignment with Bowtie 2. Nat Methods 9, 357–359.

Leone, S., Bar, D., Slabber, C.F., Dalcher, D., and Santoro, R. (2017). The RNA helicase DHX9 establishes nucleolar heterochromatin, and this activity is required for embryonic stem cell differentiation. EMBO Rep.

Li, H., and Durbin, R. (2009). Fast and accurate short read alignment with Burrows-Wheeler transform. Bioinformatics 25, 1754–1760.

Li, H., Handsaker, B., Wysoker, A., Fennell, T., Ruan, J., Homer, N., Marth, G., Abecasis, G., Durbin, R., and Genome Project Data Processing, S. (2009). The Sequence Alignment/Map format and SAMtools. Bioinformatics 25, 2078–2079.

Lu, J.Y., Shao, W., Chang, L., Yin, Y., Li, T., Zhang, H., Hong, Y., Percharde, M., Guo, L., Wu, Z., et al. (2020). Genomic Repeats Categorize Genes with Distinct Functions for Orchestrated Regulation. Cell Rep 30, 3296–3311.e3295.

Marchal, C., Sasaki, T., Vera, D., Wilson, K., Sima, J., Rivera-Mulia, J.C., Trevilla-Garcia, C., Nogues, C., Nafie, E., and Gilbert, D.M. (2018). Genome-wide analysis of replication timing by next-generation sequencing with E/L Repli-seq. Nat Protoc 13, 819–839.

Marshall, O.J., and Brand, A.H. (2015). damidseq_pipeline: an automated pipeline for processing DamID sequencing datasets. Bioinformatics 31, 3371–3373.

Mayer, C., Schmitz, K.M., Li, J., Grummt, I., and Santoro, R. (2006). Intergenic transcripts regulate the epigenetic state of rRNA genes. Mol Cell 22, 351–361.

Meshorer, E., Yellajoshula, D., George, E., Scambler, P.J., Brown, D.T., and Misteli, T. (2006). Hyperdynamic plasticity of chromatin proteins in pluripotent embryonic stem cells. Dev Cell 10, 105–116.

Misteli, T. (2011). The inner life of the genome. Sci Am 304, 66–73.

Mortazavi, A., Williams, B.A., McCue, K., Schaeffer, L., and Wold, B. (2008). Mapping and quantifying mammalian transcriptomes by RNA-Seq. Nat Methods 5, 621–628.

Nemeth, A., Conesa, A., Santoyo-Lopez, J., Medina, I., Montaner, D., Peterfia, B., Solovei, I., Cremer, T., Dopazo, J., and Langst, G. (2010). Initial genomics of the human nucleolus. PLoS Genet 6, e1000889.

Nicodemi, M., and Pombo, A. (2014). Models of chromosome structure. Curr Opin Cell Biol 28, 90–95.

Padeken, J., and Heun, P. (2014). Nucleolus and nuclear periphery: Velcro for heterochromatin. Current Opinion in Cell Biology 28, 54–60.

Peric-Hupkes, D., Meuleman, W., Pagie, L., Bruggeman, S.W., Solovei, I., Brugman, W., Graf, S., Flicek, P., Kerkhoven, R.M., van Lohuizen, M., et al. (2010). Molecular maps of the reorganization of genome-nuclear lamina interactions during differentiation. Mol Cell 38, 603–613.

Pope, B.D., Ryba, T., Dileep, V., Yue, F., Wu, W., Denas, O., Vera, D.L., Wang, Y., Hansen, R.S., Canfield, T.K., et al. (2014). Topologically associating domains are stable units of replication-timing regulation. Nature 515, 402–405.

Quinlan, A.R., and Hall, I.M. (2010). BEDTools: a flexible suite of utilities for comparing genomic features. Bioinformatics 26, 841–842.

Quinodoz, S.A., Ollikainen, N., Tabak, B., Palla, A., Schmidt, J.M., Detmar, E., Lai, M.M., Shishkin, A.A., Bhat, P., Takei, Y., et al. (2018). Higher-Order Inter-chromosomal Hubs Shape 3D Genome Organization in the Nucleus. Cell 174, 744–757 e724.

Ragoczy, T., Telling, A., Scalzo, D., Kooperberg, C., and Groudine, M. (2014). Functional redundancy in the nuclear compartmentalization of the late-replicating genome. Nucleus 5, 626–635.

Ramirez, F., Dundar, F., Diehl, S., Gruning, B.A., and Manke, T. (2014). deepTools: a flexible platform for exploring deep-sequencing data. Nucleic Acids Res 42, W187–191.

Ran, F.A., Hsu, P.D., Wright, J., Agarwala, V., Scott, D.A., and Zhang, F. (2013). Genome engineering using the CRISPR-Cas9 system. Nature protocols 8, 2281–2308.

Reynolds, R.C., Montgomery, P.O., and Hughes, B. (1964). Nucleolar “Caps” Produced by Actinomycin D. Cancer Res 24, 1269–1277.

Robinson, J.T., Thorvaldsdóttir, H., Winckler, W., Guttman, M., Lander, E.S., Getz, G., and Mesirov, J.P. (2011). Integrative genomics viewer. Nature Biotechnology.

Sampath, P., Pritchard, D.K., Pabon, L., Reinecke, H., Schwartz, S.M., Morris, D.R., and Murry, C.E. (2008). A hierarchical network controls protein translation during murine embryonic stem cell self-renewal and differentiation. Cell Stem Cell 2, 448–460.

Savic, N., Bar, D., Leone, S., Frommel, S.C., Weber, F.A., Vollenweider, E., Ferrari, E., Ziegler, U., Kaech, A., Shakhova, O., et al. (2014). lncRNA Maturation to Initiate Heterochromatin Formation in the Nucleolus Is Required for Exit from Pluripotency in ESCs. Cell Stem Cell 15, 720–734.

Savić, N., Bär, D., Leone, S., Frommel, Sandra C., Weber, Fabienne A., Vollenweider, E., Ferrari, E., Ziegler, U., Kaech, A., Shakhova, O., et al. (2014). lncRNA Maturation to Initiate Heterochromatin Formation in the Nucleolus Is Required for Exit from Pluripotency in ESCs. Cell Stem Cell 15, 720–734.

Stansfield, J.C., Cresswell, K.G., Vladimirov, V.I., and Dozmorov, M.G. (2018). HiCcompare: an R-package for joint normalization and comparison of HI-C datasets. BMC Bioinformatics 19, 279.

Sullivan, G.J., Bridger, J.M., Cuthbert, A.P., Newbold, R.F., Bickmore, W.A., and McStay, B. (2001). Human acrocentric chromosomes with transcriptionally silent nucleolar organizer regions associate with nucleoli. EMBO J 20, 2867–2874.

van Koningsbruggen, S., Gierlinski, M., Schofield, P., Martin, D., Barton, G.J., Ariyurek, Y., den Dunnen, J.T., and Lamond, A.I. (2010). High-resolution whole-genome sequencing reveals that specific chromatin domains from most human chromosomes associate with nucleoli. Mol Biol Cell 21, 3735–3748.

van Steensel, B., and Belmont, A.S. (2017). Lamina-Associated Domains: Links with Chromosome Architecture, Heterochromatin, and Gene Repression. Cell 169, 780–791.

van Steensel, B., Delrow, J., and Henikoff, S. (2001). Chromatin profiling using targeted DNA adenine methyltransferase. Nat Genet 27, 304–308.

Vertii, A., Ou, J., Yu, J., Yan, A., Pages, H., Liu, H., Zhu, L.J., and Kaufman, P.D. (2019). Two contrasting classes of nucleolus-associated domains in mouse fibroblast heterochromatin. Genome Res 29, 1235–1249.

Weeks, S.E., Metge, B.J., and Samant, R.S. (2019). The nucleolus: a central response hub for the stressors that drive cancer progression. Cell Mol Life Sci 76, 4511–4524.

Yu, G., Wang, L.G., and He, Q.Y. (2015). ChIPseeker: an R/Bioconductor package for ChIP peak annotation, comparison and visualization. Bioinformatics 31, 2382–2383.

